# Loss of Kallmann syndrome-associated gene WDR11 disrupts primordial germ cell development by affecting canonical and non-canonical Hedgehog signalling

**DOI:** 10.1101/2020.09.06.284927

**Authors:** Jiyoung Lee, Yeonjoo Kim, Paris Ataliotis, Hyung-Goo Kim, Dae-Won Kim, Dorothy C. Bennett, Nigel A. Brown, Lawrence C. Layman, Soo-Hyun Kim

## Abstract

Mutations of *WDR11* are associated with Kallmann syndrome (KS) and congenital hypogonadotrophic hypogonadism (CHH), typically caused by defective functions of gonadotrophin-releasing hormone (GnRH) neurones in the brain. We previously reported that Wdr11 knockout mice show profound infertility with significantly fewer germ cells present in the gonads. To understand the underlying mechanisms mediated by WDR11 in these processes, we investigated the effects of *Wdr11* deletion on primordial germ cell (PGC) development. Using live-tracking of PGCs and primary co-cultures of genital ridges (GR), we demonstrated that *Wdr11*-deficient embryos contained reduced numbers of PGCs which had delayed migration due to significantly decreased proliferation and motility. We found primary cilia-dependent canonical Hedgehog (Hh) signalling was required for proliferation of the somatic mesenchymal cells of GR, while primary cilia-independent non-canonical Hh signalling mediated by Ptch2/Gas1 and downstream effectors Src and Creb was required for PGC proliferation and migration, which was disrupted by the loss of function mutations of WDR11. Therefore, canonical and non-canonical Hh signalling are differentially involved in the development of somatic and germ cell components of the gonads, and WDR11 is required for both of these pathways operating in parallel in GR and PGCs, respectively, during normal PGC development. Our study provides a mechanistic link between the development of GnRH neurones and germ cells mediated by WDR11, which may underlie some cases of KS/CHH and ciliopathies.

## INTRODUCTION

Genetic defects affecting the development, integration and coordination of the hypothalamic-pituitary-gonadal (HPG) axis constitute most of the aetiologies in Kallmann syndrome (KS) and idiopathic congenital hypogonadotrophic hypogonadism (CHH), clinically defined by low plasma levels of sex steroids and gonadotropins with absent/delayed sexual maturation and infertility. KS patients also present with anosmia, a lack of the sense of smell (1). Current doctrine is that CHH/KS is a hypothalamic and/or pituitary disease caused by inappropriate development or failed reactivation of gonadotrophin-releasing hormone (GnRH) neurons at puberty. Therefore, infertility in patients with KS/CHH is routinely treated by GnRH or gonadotropin replacement therapy (2;3). The majority of male patients (75-95%) show normalised testosterone levels after treatment. However, only 5-20% of them achieve normal sperm concentrations and 20-40% show azoospermia, while the remainder exhibit severe oligospermia (1;4–6). These findings suggest that primary defects in the gonads may exist in these individuals.

Previously, we identified *WDR11* as a genetic locus for KS/CHH (7). Missense variants of *WDR11* have also been reported in septo-optic dysplasia, combined pituitary hormone deficiency and pituitary stalk interruption syndrome (8–10). WDR11 belongs to a family of proteins with the evolutionarily conserved WD40-repeat (WDR) domains, forming β-propeller structures known to mediate protein-protein interactions (7;11). Our previous studies of a Wdr11 knockout (KO) mouse model have indicated a critical role for *WDR11* in development (12). Wdr11 is required for normal ciliogenesis as loss of Wdr11 resulted in short and infrequent primary cilia. Since multiple developmental signalling pathways functionally rely on primary cilia, the majority of Wdr11 KO embryos die in utero at mid-gestation (after E12.5) with severe developmental defects (12). Those rare individual mice that survived through adulthood display features overlapping with KS/CHH such as delayed puberty and infertility, accompanied by reduced levels of GnRH and gonadotrophins. Migration of GnRH neurones is disrupted in these mice causing reduced total numbers of GnRH neurones reaching the hypothalamus. In addition, Wdr11 KO mice are born with hypoplastic gonads containing fewer germ cells compared to the wild type (WT) littermates. Wdr11-deficient testes are smaller in size and contain fewer spermatocytes and spermatids with an increased frequency of morphologically abnormal sperm found in the seminiferous tubules (12). Wdr11-deficient ovaries are also smaller than WT and present with disproportionally higher numbers of oogonia or primordial follicles and reduced numbers of mature follicles (12). These data indicate that loss of Wdr11 results in defective development of germ cells and gonads in both sexes.

Primordial germ cells (PGCs) are bipotential stem cells and the founders of gametes. They undergo distinctive developmental stages (specification, polarization, migration and invasion) before they become immobile and differentiate into either spermatozoa or oocytes in the gonads (13–15). In mouse, PGCs originate from the posterior primitive streak (E7.5) and move into the developing hindgut where they migrate along its anterior extension (E8-E9.5). Then they move out of the hindgut, travel through the mesentery of the dorsal body wall, and finally enter the bilateral genital ridges (GR) (E10.5) (13;14). Normal development and migration of PGCs are regulated by networks of signalling molecules and receptors expressed in the microenvironment of the germ cell niche, including chemokine SDF1 and its receptor CXCR4 (16;17). Interestingly, SDF1 and CXCR4 are also important in GnRH neuron migration, and decreased numbers of GnRH neurons are observed in *Cxcr4* knockout mice (18).

Hedgehog (Hh) is a major morphogen that binds to the cell surface receptors Ptch1 and Ptch2. Boc (bioregional Cdon binding protein), Cdon (cell-adhesion-molecule-related/downregulated by oncogenes, also called as Cdo) and Gas1 (growth arrest-specific gene 1) are membrane-associated co-receptors that interact with the primary Ptch receptors (19–22). They can bind Hh ligand independently of Ptch, facilitating ligand–receptor interactions at the cell surface (19). We recently reported that they are critically required for selective activation of Smo-downstream signalling (20). Gli transcription factors mediate canonical Hh signalling pathway via primary cilia-dependent mechanisms. Gli-independent non-canonical signalling also occurs, which does not require Smo localisation to the primary cilia (20;23;24). Studies in Drosophila and zebrafish have previously demonstrated that Hh signalling is involved in the development of PGCs, but not as a guidance cue or fate determinant (25–27). It was shown that Sonic Hedgehog (Shh) is not a chemo-attractant for PGCs in the mouse but non-canonical Hh signalling mediated by the Ptch2/Gas1 receptor complex, exclusively expressed on the surface of PGCs, is important in PGC motility (20). Notably, putative mutations of Hh signalling pathway genes including GLI, SMO and PTCH1 have been suggested in KS/CHH (28–30).

Here we investigate the potential involvement of the KS/CHH-associated gene WDR11 in the establishment of germ cells in the gonads. Our data demonstrate that Wdr11 is essential for the proliferation and migration of PGCs as well as the growth of the surrounding soma, mediated by non-canonical and canonical Hh signalling, respectively. We propose that in addition to the hypothalamic GnRH deficiency, primary defects in the germ cells may underlie KS/CHH patients with WDR11 mutations. The mechanisms we revealed may apply to other ciliopathies where hypogonadism and infertility are part of the clinical features.

## RESULTS

### WDR11 is expressed in the PGC developmental niche

Initial analyses by RT-PCR showed that Wdr11 was expressed in the developing and adult urogenital organs of both sexes. Wdr11 mRNA was present in the regions through which PGCs migrate including the hindgut (HG) at E9.5 and the urogenital ridge area (UG) at E10.5 - 11.5. Wdr11 was also expressed in the post-pubertal testis, epididymis, ovary and kidney (Fig 1A). The spatio-temporal expression of Wdr11 in the developing urogenital system was further demonstrated by whole mount X-gal staining in Wdr11 heterozygote embryos (Fig 1C) and by direct immunofluorescence staining for Wdr11 (Fig 1D). Both methods indicated widespread expression in gonadal development including HG and mesonephric tubules. Co-immunostaining with SSEA1, a carbohydrate antigen specifically expressed by PGCs, confirmed the expression of Wdr11 in individual PGCs as well as in the surrounding somatic cells, showing diffuse peri-nuclear and cytoplasmic signal, which was absent in Wdr11 KO embryos (Fig 1D). Based on this broad expression of Wdr11 in both the mesenchymal and germ cell components of the gonads, we hypothesised that Wdr11 may have broad effects in reproductive system development including PGCs.

**Figure 1.**
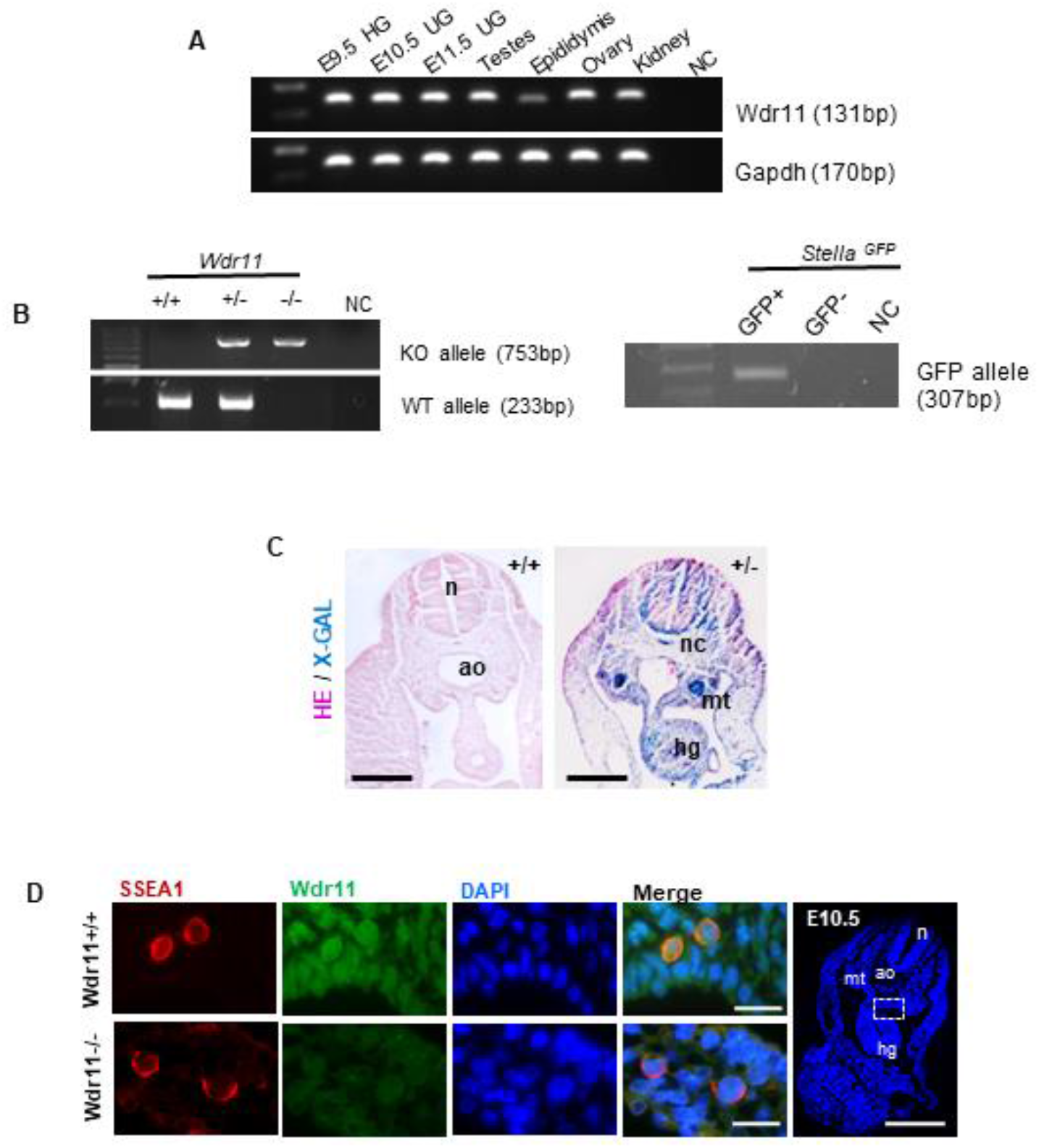
Wdr11 is expressed in embryonic urogenital tissues and PGCs. **(A)** Expression of Wdr11 and Gapdh in the hindgut (HG) at E9.5, the urogenital ridge area (UG) at E10.5 - 11.5 and the post-pubertal reproductive organs (testis, epididymis, ovary) and kidney in 8-week-old mice was assessed by RT-PCR. Representative results are shown from 3 independent biological samples. **(B)** Genotyping analyses of Wdr11 knockout mice by PCR. The WT and gene-trap allele are indicated (left panel). The presence of the GFP-specific allele in the *StellaGFP;Wdr11* hybrid line was confirmed by PCR (right panel) as well as by a test breeding (see Materials and Methods). **(C)** Images of transverse sections of whole-mount X-gal-stained paraffin-embedded E10.5 embryos with eosin counterstaining. WT (+/+) and heterozygous (+/−) embryos are shown (scale bar, 100μm). n, neural tube; ao, aorta; nc, notochord; mt, mesonephric tubules; hg, hindgut. **(D)** Immunofluorescence staining of WDR11 (green) and DAPI (blue) on the transverse sections of Wdr11 WT and KO embryos at E10.5. Scale bar represents 100μm. **(E)** Transverse sections of E10.5 *Wdr11* WT and KO embryos stained with anti-WDR11 (green), anti-SSEA1 (red) and DAPI (blue), confirming expression of Wdr11 in PGCs (SSEA1-positive) and the surrounding somatic cells in the GR area. A zoomed-in image of the dotted area is shown. Scale bar represents 20μm.

### Loss of Wdr11 affects PGC migration

We have previously reported that the gonads of Wdr11-deficient mice were unusually small, which may be caused, at least in part, by a deficiency of germ cells (12). Wdr11 deficient mice showed ovaries containing reduced numbers of oocytes and testes containing significantly fewer spermatocytes (12). Since defective development of PGCs during early embryogenesis can result in insufficient numbers of germ cells present in the gonads at birth, leading to in/sub-fertility or premature ovarian failure, we investigated whether the absence of Wdr11 has any impact on PGC development.

First, we analysed the number and location of PGCs by anti-SSEA1 immunofluorescence and alkaline phosphatase staining at different developmental stages (E9.5 – E11.5). The results showed that Wdr11-null embryos were still populated with PGCs in their normal migratory path between the HG and GR, but many of them were inappropriately located for the stage of development (Fig 2A). Therefore, loss of Wdr11 did not completely prevent the specification of PGCs, but disrupted their migration. When we quantified the total numbers of PGCs by counting the SSEA1-positive cells, there was a significant reduction in Wdr11-/- embryos compared to WT (Fig 2B). Our analyses confirmed an inappropriate accumulation of PGCs in the hindgut and mesentery, compared to WT at E10.5 (Fig 2C). This led to a significantly lower number of PGCs arriving in the GR at E10.5 in Wdr11-/- embryos (Fig 2C), and concomitantly a significantly higher number of ectopic PGCs (Fig 2D). Mis-localised PGCs fail to develop normally owing to the lack of survival signals from their environment. These data suggest that Wdr11 KO caused impaired migration of PGCs, affecting germ cell establishment in the future gonads.

**Figure 2.**
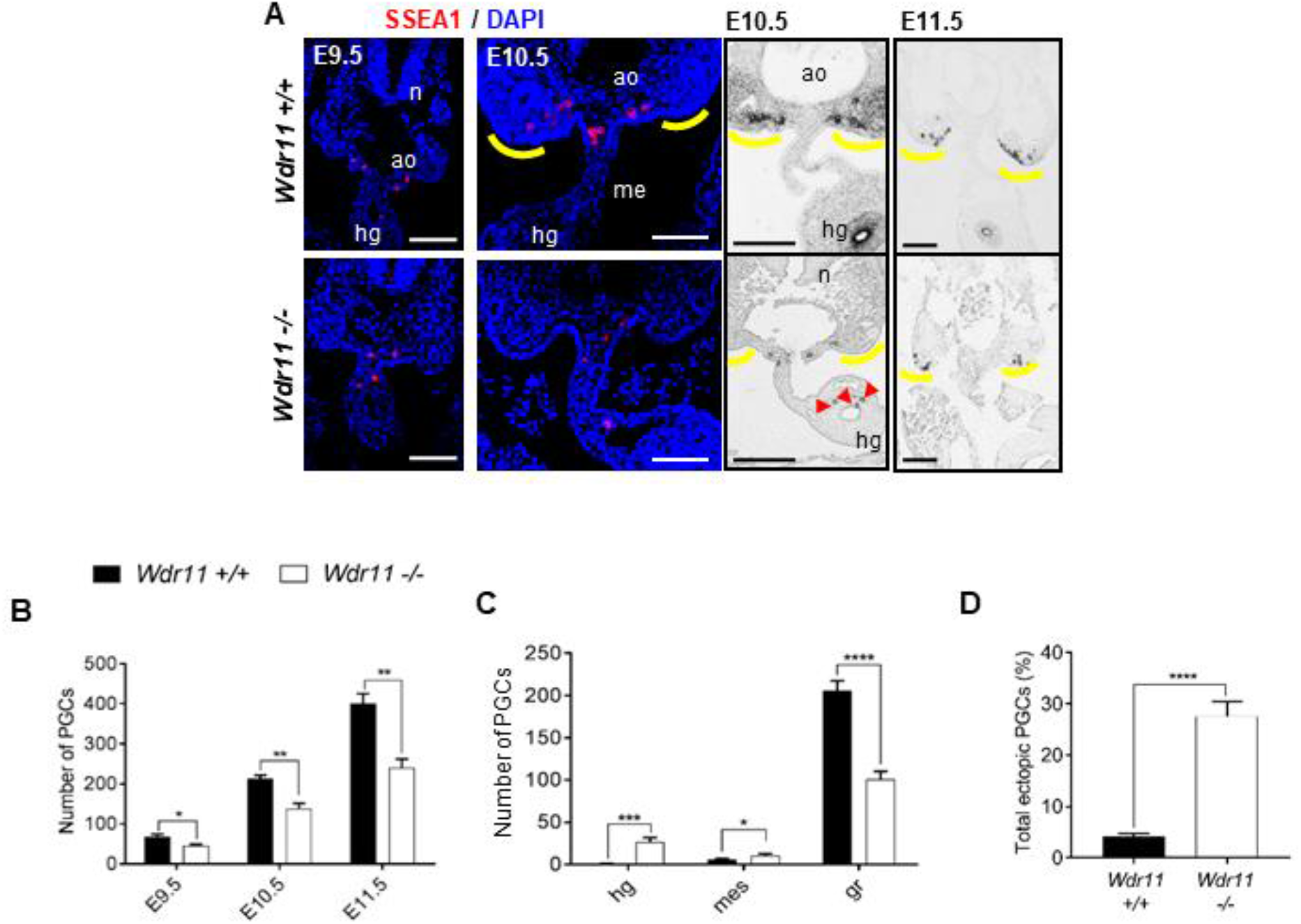
Wdr11 KO causes a reduction in total PGCs with increased ectopic PGCs. **(A)** Immunofluorescence (left) and alkaline phosphatase (right) staining analyses of PGCs in their migratory route from hindgut towards the GRs (yellow outlines) at different developmental stages as indicated. Compared to the WT, Wdr11-/- embryos contain fewer PGCs in the GRs, and more ectopically located PGCs (red arrowheads). n, neural tube; ao, aorta; hg, hindgut; me, mesentery. Scale bar, (E9.5) 20μm; 100μm (E10,5, E11.5). **(B)** Wdr11 KO embryos show a reduction in total PGC numbers. Total PGCs were counted from every other slide of the serial sections of E9.5, E10.5 and E11.5 embryos (n=5 per genotype). Total count per embryo is shown as mean ± SD. **(C)** Wdr11 KO embryos contain an increased number of ectopic PGCs. Total number of PGC population in each location at E10.5 are shown for each genotype (n=5). hg, hindgut; mes, mesentery; gr, genital ridge. **(D)** The proportion of ectopic PGCs present in WT and KO embryos (n=5 per genotype) at E10.5 is shown as a percentage value of total PGCs. Error bars represent SD. Unpaired Student’s t-test (*P < 0.05; **P < 0.01; ***P < 0.001; ****P < 0.0001).

### Defective motility of Wdr11-deficient PGCs

We hypothesised that Wdr11 KO would affect at least one of the following key processes in PGC development: motility, proliferation or survival. First, we examined the motile behaviours of PGCs and quantitatively analysed their intrinsic motility. For this purpose, we have established embryo slice cultures of explanted GR of mice expressing a Stella promoter-driven GFP transgene (*Stella*^*GFP*^). Stella is the most specific marker for PGCs, being expressed soon after their specification at ~E7.5 and maintained until E13.5 in females and E15.5 in males (31). We performed a time-lapse live imaging and motion analysis of PGCs in WT and Wdr11 KO background using a *Stella*^*GFP*^;*Wdr11* hybrid strain at E10.5. The movements of PGCs in the time-lapse movies were manually tracked and analysed for directionality, targeting, distance and speed. When the migration over time (>10 hours) was compared in WT and Wdr11-null embryos, we found that PGCs in both genotypes were moving towards the GR area, showing no discernible differences in their targeting and directionality of migration (Fig 3A). To examine if Wdr11 deficiency altered the intrinsic motility of PGCs, we performed quantitative motion analyses, which revealed that the velocity, accumulated distance and Euclidean distance of migration were significantly reduced in the Wdr11 KO embryos (Fig 3B), consistent with the ectopic distribution of PGCs observed in our immunofluorescence experiments (Fig 2). Directionality values representing the degree to which the migratory path of a cell strayed from a straight line were not altered. Hence, the majority of the cells were still moving towards the GRs but the speed and distance of migration were reduced in Wdr11 KO, resulting in a significantly reduced number of PGCs arriving at the GRs.

**Figure 3.**
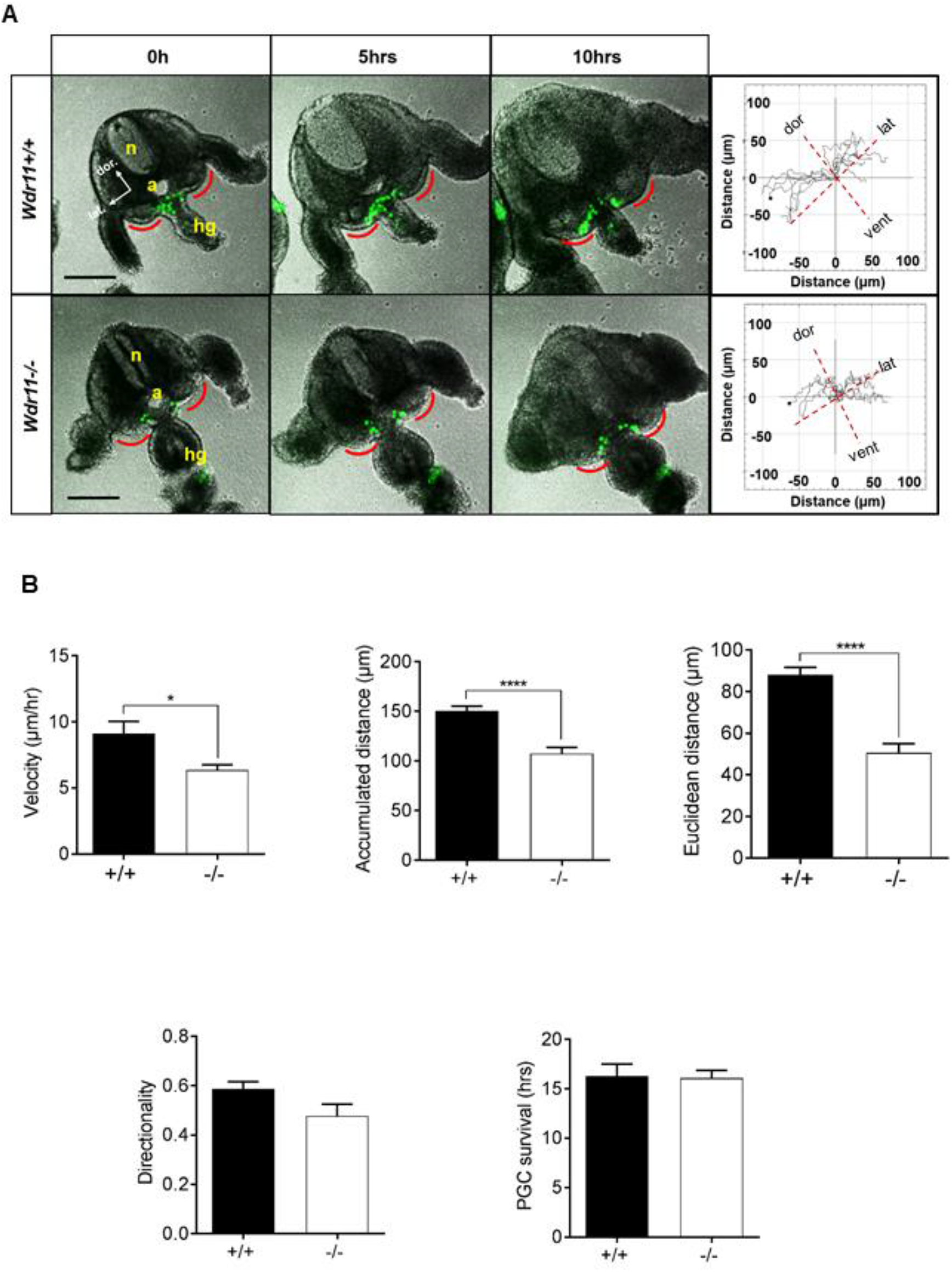
Loss of *Wdr11* disrupts PGC migration. **(A)** Time-lapse images of embryo slice cultures from *Stella*^*GFP*^;*Wdr11* hybrid mice. Representative images of a plane of the z-stack at 0, 5 and 10 hours are shown from movies of E10.5 *Stella*^*GFP+/+*^;*Wdr11*^*+/+*^ and *Stella*^*GFP+/+*^;*Wdr1*^*-/-*^ embryos in biologically independent experiments (Supplementary Movies 1 and 2). n, neural tube; a, aorta; hg; hindgut. Scale bar represents 100μm. The PGCs (GFP-positive) migrating towards the GRs (indicated in red) were tracked using ImageJ. Trajectory plots of migration (right panel) were generated by placing the starting points of individual PGC tracks onto the same point. Origins of all tracks were centred at the 0,0 XY coordinate with distance in micrometers on x- and y-axes. The direction of migration in relevance to the embryo orientation is shown with dotted lines. Dor, dorsal; lat, lateral; vent, ventral. **(B)** Comparison of the velocity (total accumulated distance over time period), accumulated distance (total path travelled) and Euclidean distance (the shortest distance between the start and end points) indicates a significant reduction in PGC migration in Wdr11-null embryos compared to WT. There was no significant difference between the two genotypes in the directionality of migration (the straightness of the migration path, represented by the ratio between Euclidean distance and accumulated distance) or cell survival (assessed by the number of hours that the GFP fluorescence from a cell was detected during the imaging shown in A). Error bars represents SEM. Data were from independent slice cultures (WT, n=9; KO, n=6) where 7-10 PGCs were tracked from each embryo slice. Statistical analysis by unpaired Student’s t-test (*P < 0.05; ****P < 0.0001; directionality, *P* = 0.061; survival, *P* = 0.64).

We observed some PGCs disintegrating progressively in our live-imaging, a hallmark of apoptotic cells (see supplementary movies 1 and 2). To ascertain whether the reduction in movement was due to reduced survival of PGCs in the mutants, we assessed the survival times by measuring the mean number of hours that the green fluorescence of individual cells could be observed in the movies and found no difference between the genotypes (Fig 3B). Combined, these data suggest that Wdr11 is necessary for active PGC motility during migration towards the GRs but not the targeting and attraction of PGCs towards GRs, nor in maintaining survival.

### Defective proliferation but normal apoptosis in Wdr11 mutants

One possible explanation for the decreased number of PGCs is reduced proliferation. In mice, PGCs continue to proliferate during and after migration, rapidly expanding to a final population of ~25,000 cells per embryo at E13.5. Indeed, PGCs visibly divided during time-lapse imaging (see supplementary movies 1 and 2). To determine if loss of Wdr11 affects PGC proliferation, we quantitatively assessed mitotically active cells by phosphorylated-histone H3 (PH3) staining of E10.5 GR (Fig 4A). The total number of PH3-positive and -negative PGCs and the mesenchymal somatic cells (GFP-negative, DAPI-positive) between the forelimb and hindlimb buds were manually counted and PH3-labelling index values generated. Wdr11-/- embryos showed a lower PH3 labelling index compared to WT in both PGCs and mesenchymal cells (Fig 4B). Therefore, Wdr11 is required for the proliferation of both PGCs and mesenchymal cells in the migratory niche.

**Figure 4.**
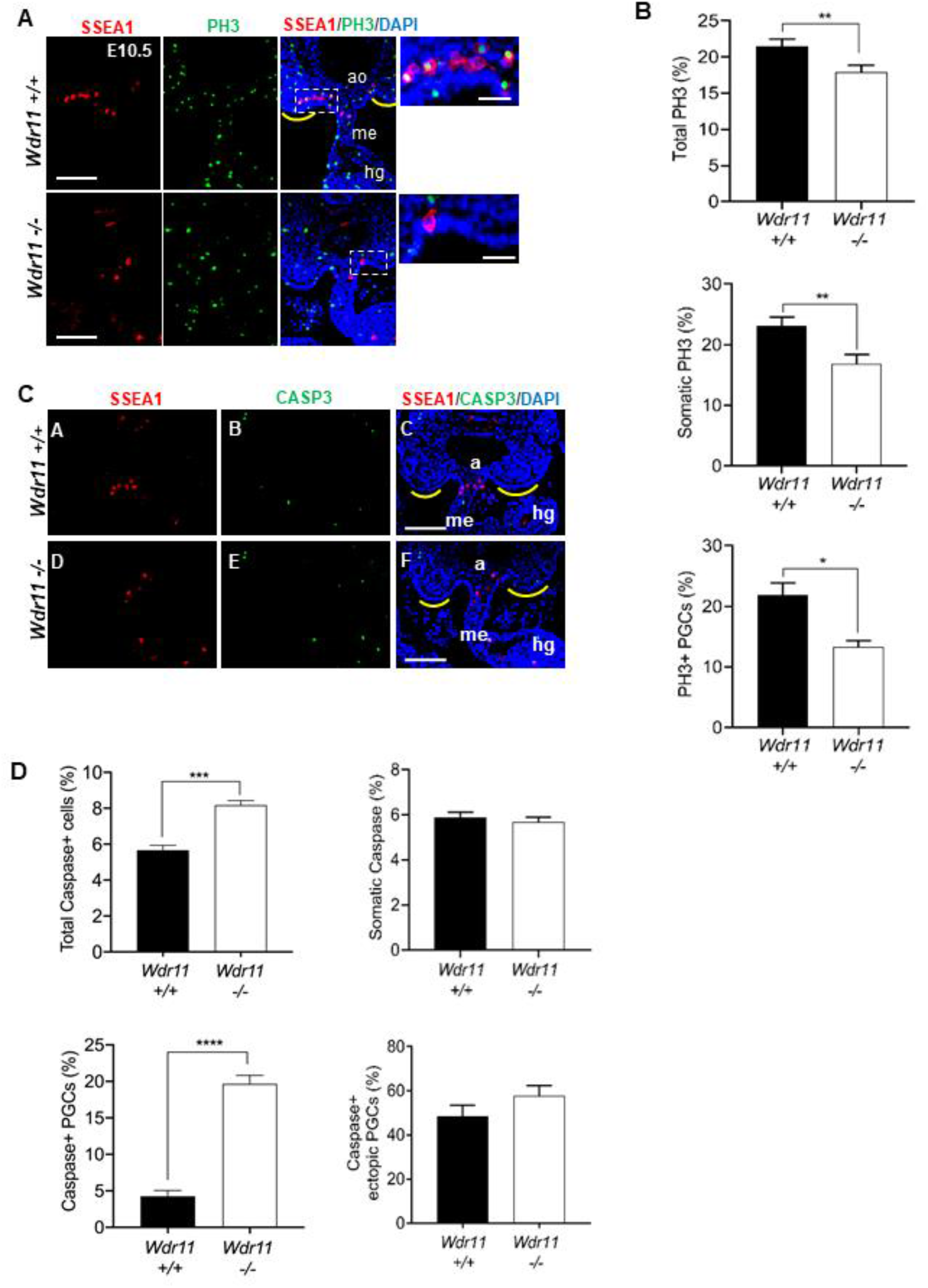
*Wdr11* KO affects PGC proliferation but not directly their survival. **(A)** Representative images of immunofluorescence staining of SSEA1 (red) and phospho-histone H3 (PH3, green) on Wdr11 WT and KO embryos at E10.5. Yellow line indicates the GRs. Zoomed-in images of the dotted area are shown on the right. ao, aorta; hg, hindgut; me, mesentery. Scale bar, 100μm (left); 20μm(right). **(B)** Analyses of cell proliferation based on the PH3-positive cell counts. The percentage values are obtained by manually counting the total positive cells against the total cell counts labelled with DAPI from every other section of the PGC migratory route (top panel). The percentage of PH3-positive cells in the somatic cell population (middle panel) and the PGC population (bottom panel) are compared between WT and KO embryos. Error bars represent SEM. Statistical analysis by unpaired Student's t-test (n=5, number of embryos for each genotype; *P < 0.05; **P < 0.01). **(C)** Representative images of immunofluorescence staining of SSEA1 (red) and Caspase-3 (green) on E10.5 Wdr11 WT and KO embryos. Yellow line indicates the GRs. Scale bar, 100μm. **(D)** Analyses of apoptosis based on Caspase-3 immunofluorescence. CASP3-positive and total cells were counted from every other section of the PGC migratory route (top left). The percentages of CASP3-positive somatic cells (top right), PGCs (bottom left) and ectopic PGCs (bottom right) are compared between WT and KO embryos. Error bars represent SEM. Statistical analysis by unpaired Student's t-test (n=5, number of embryos for each genotype; ***P<0.001).).

During PGC migration, there is an up-regulation of factors involved in apoptosis, and embryos with a defective apoptotic pathway exhibited ectopic PGCs that were not cleared effectively (32). To determine further whether loss of Wdr11 altered apoptosis, we carried out immunostaining for cleaved-Caspase 3 and manually counted Casp3-positive and - negative cells against total cell counts. The results indicated a significant increase of total apoptotic cells in Wdr11 KO embryos. However, upon careful examination, we found that this was due to the abnormally elevated numbers of ectopic PGCs present in these embryos, rather than enhanced apoptosis in general. This conclusion was based on the fact that the apoptotic index in the mesenchymal somatic cells was not different between the genotypes and the ectopic PGCs were equally positive for Casp3 in both WT and Wdr11 KO (Fig 4B). Therefore, loss of Wdr11 did not result in an overall increase in cell death, confirming our observation from the time-lapse imaging (Fig 3B).

### PGC developmental signalling in Wdr11 mutants

It is possible that the defective establishment of PGCs in embryos lacking Wdr11 simply reflects an overall retardation in development. To exclude such a notion, we validated the developmental stages of the embryos used in our analyses by morphological landmarks such as somite numbers, absence/presence of hind limbs and tail buds (E9.5 and E10.5, respectively) and the closure of lens vesicle (E11.5), which indicated that mutant embryos did not have general developmental defects, at least during the period we studied.

We next examined whether Wdr11 KO affected the expression of genes known to play critical roles in the development of PGCs such as Blimp1, c-Kit, Steel (Kitl), Cxcr4 and Sdf1 (Cxcl12). Our initial screening confirmed the expression of these genes in the WT PGC migratory niche and adult urogenital organs (Fig 5A). Quantitative analyses by RT-qPCR indicated that Wdr11-/- embryos did not show significantly reduced mRNA levels of these regulators, except for a significant decrease in c-Kit (Fig 5B). There was also a numerical but non-significant reduction in Cxcr4. This is consistent with the reduced total number of PGCs in the mutants, as both c-Kit and Cxcr4 are cell surface receptors expressed by PGCs, while Cxcr4 is also more widely expressed (33;34). The expression of the respective ligands for these receptors, Steel and Sdf1, which mediate the chemo-attraction of PGCs towards the gonads, was not altered (Fig 5B). Therefore, the reduction in c-Kit alone seemed unlikely to explain the decreased proliferation of PGCs and mesenchymal somatic cells, nor the reduced PGC migration in Wdr11−/− embryos.

**Figure 5.**
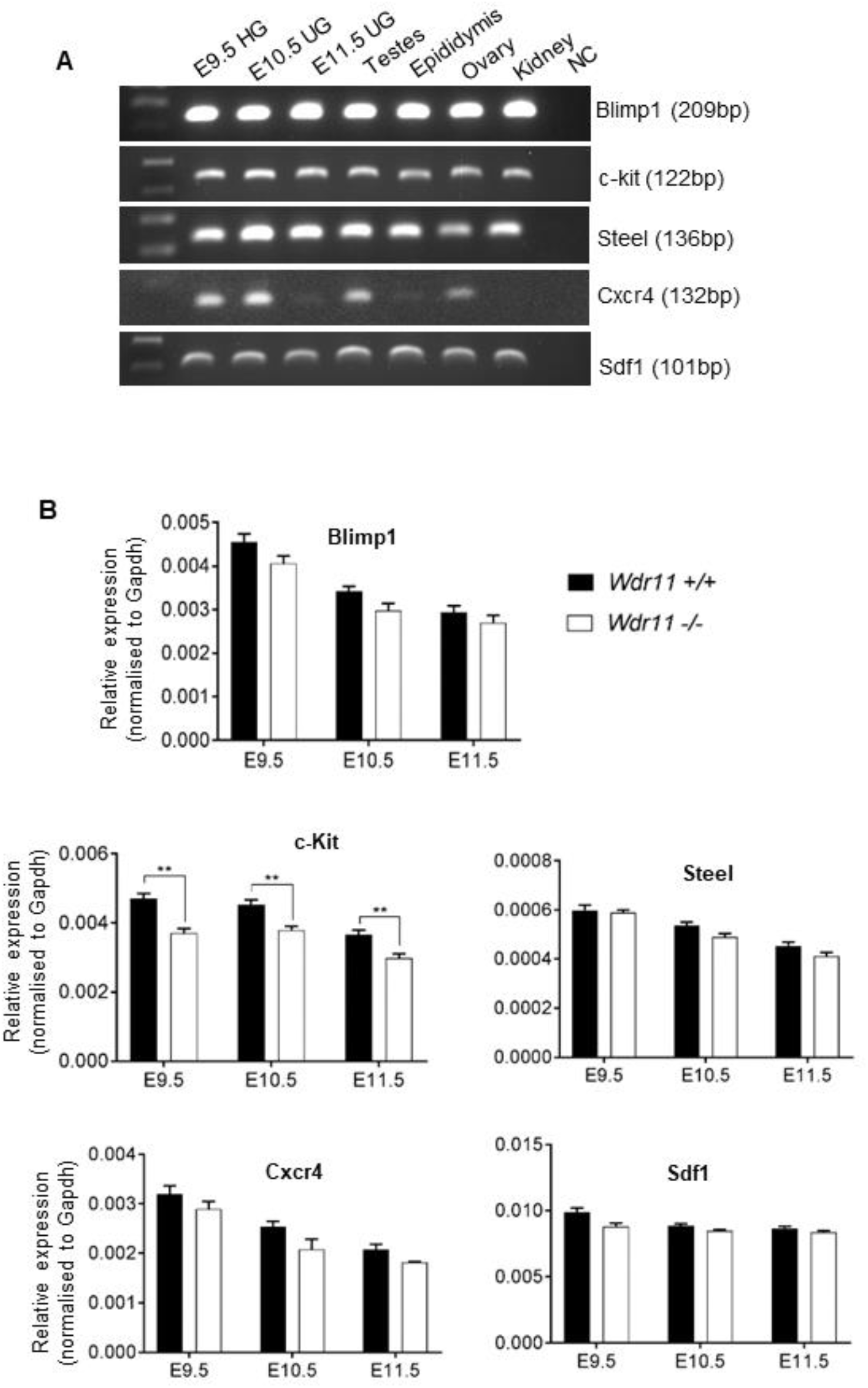
Wdr11 KO did not affect the expression of key genes required for the PGC migration and urogenital development. **(A)** RT-PCR analysis of Blimp1, c-Kit, Steel, Cxcr4 and Sdf1 mRNA in the PGC migratory niche of E9.5, E10.5 and E11.5 embryos (HG, hindgut; UG, urogenital ridge) and the reproductive organs and kidney in adults. NC, no template control. **(B)** Comparison of gene expression in WT and Wdr11 KO by qRT-PCR. Loss of Wdr11 did not affect expression of these genes in the PGC migratory niche area, except c-kit showed a decrease. Values are shown as means ± SD. Statistical analysis by unpaired Student's t-test (n=5 embryos for each genotype; **P < 0.01).

### Wdr11 KO disrupts primary cilia and canonical Hh signalling

Previously we reported that Wdr11 is required for ciliogenesis and cells lacking Wdr11 display short and infrequent primary cilia (12). Given the important role of primary cilia in developmental signalling such as the Hh pathway, we hypothesised that the PGC deficiency in Wdr11 KO mice may be due to disruption in cilia-dependent signalling in the GR regions. Notably, the pluripotent PGCs are naturally un-ciliated but remain responsive to Hh signalling (20). Immunofluorescence staining of GR sections for ciliary marker Arl13b confirmed that the mesenchymal cells immediately surrounding the PGCs were ubiquitously ciliated in WT (Fig 6A and C). The mesenchymal cells in Wdr11 KO embryos, however, displayed significantly short and fewer cilia (Fig 6A and B). We have previously demonstrated that Hh signalling pathway genes are expressed in the PGC migratory niche in mice (20). Since Hh signalling is known to regulate proliferation of different cell types (35–37), we investigated if the reduced somatic cell proliferation in Wdr11 mutants was linked to an attenuation of Hh signalling caused by defective cilia. We first examined the expression of Ptch1 and Gli1/2/3 in the PGC migration routes in WT embryos, which demonstrated a significant induction of these genes from E9.5, reaching a maximum at E10.5 followed by a gradual decrease till E12.5 (Fig 7A). These results suggest active canonical Hh signalling in this site and period. Our analyses of Wdr11-null GR, however, showed significantly diminished expression of these genes even at the E10.5 peak (Fig 7B), suggesting a severe defect in canonical Hh signalling in the absence of Wdr11. Boc is an obligatory co-receptor for Ptch1 and mediates de-repression of Smo upon Hh ligand reception by Ptch1 (19;20;38). We have previously shown that Boc is broadly expressed in the PGC migratory niche on both somatic cells and PGCs (20). When we assessed Boc expression in Wdr11-null GR, significant reductions in both mRNA (Fig 7C) and protein levels (Fig 8B) were observed. Therefore, general depression of the canonical Hh signalling involving insufficient expression of Ptch1 and Boc may underlie the defective mesenchymal cell proliferation in Wdr11 KO.

**Figure 6.**
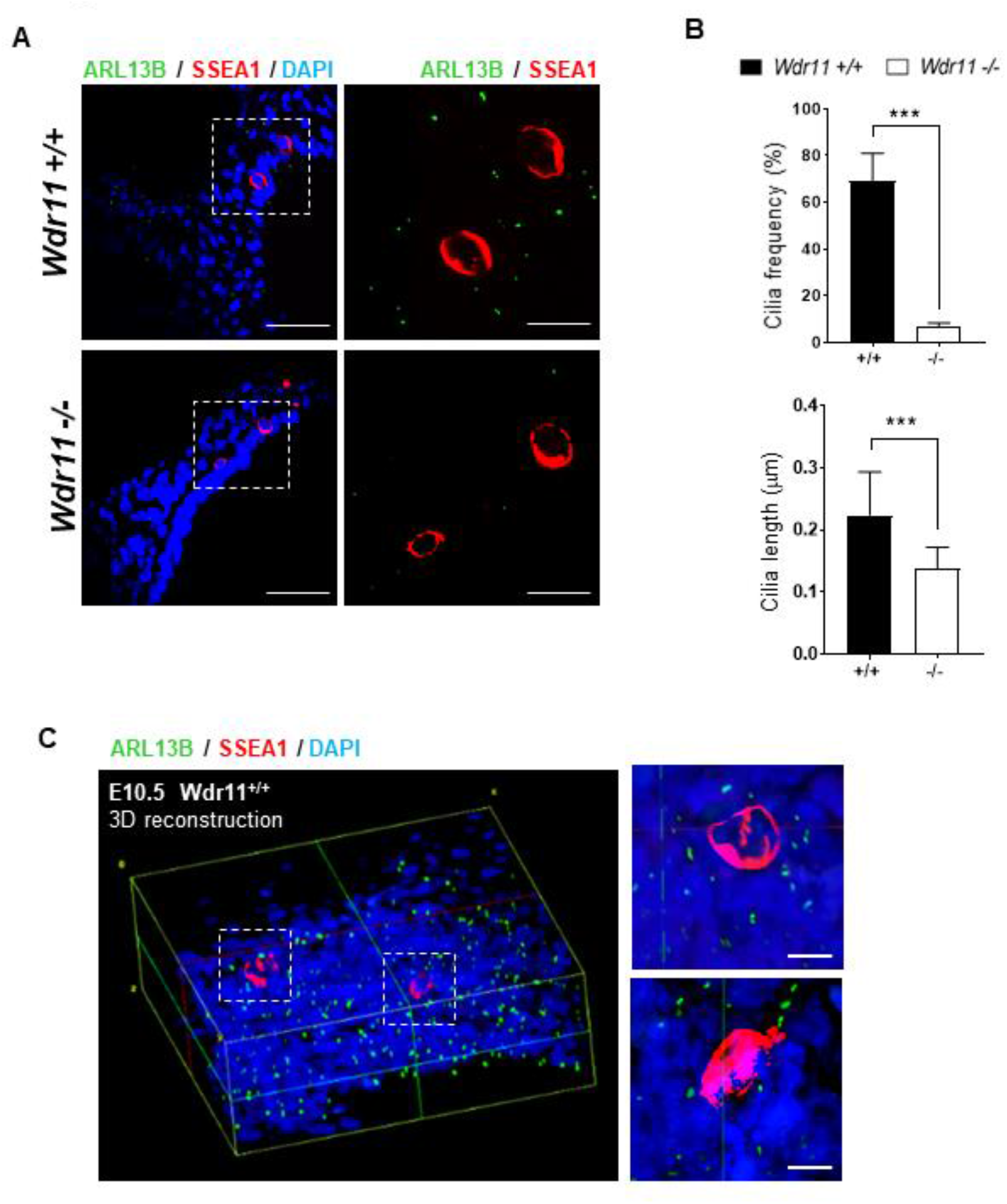
Primary cilia are disrupted in the Wdr11 KO PGC migratory niche. **(A)** Immunofluorescence analyses of primary cilia by anti-ARL13B (green) and anti-SSEA1 (red) staining on GR tissue sections of Wdr11 WT and KO embryos at E10.5. Zoomed-in images of the dotted area are shown on the right. Scale bar, 10 μm (left panels) and 50 μm (right panels). Representative images are shown from 5 independent biological samples. **(B)** Comparison of the ciliation frequency and cilium length observed in GR sections. Ciliation frequency values are generated from the total number of cilia and nuclei counted from the maximum intensity projection images of each channel manually. The length of cilia was assessed by measuring the maximum projection of Arl13b signal using ImageJ. WT (n=119) and KO (n=103). Unpaired t-test. ***P < 0.001. **(C)** 3D reconstruction of GR region in E10.5 WT embryo stained with anti-Arl13B and anti-SSEA1. ImageJ software using the volume viewer plugin was used to build the image stacks. Scale bar, 10 μm.

**Figure 7.**
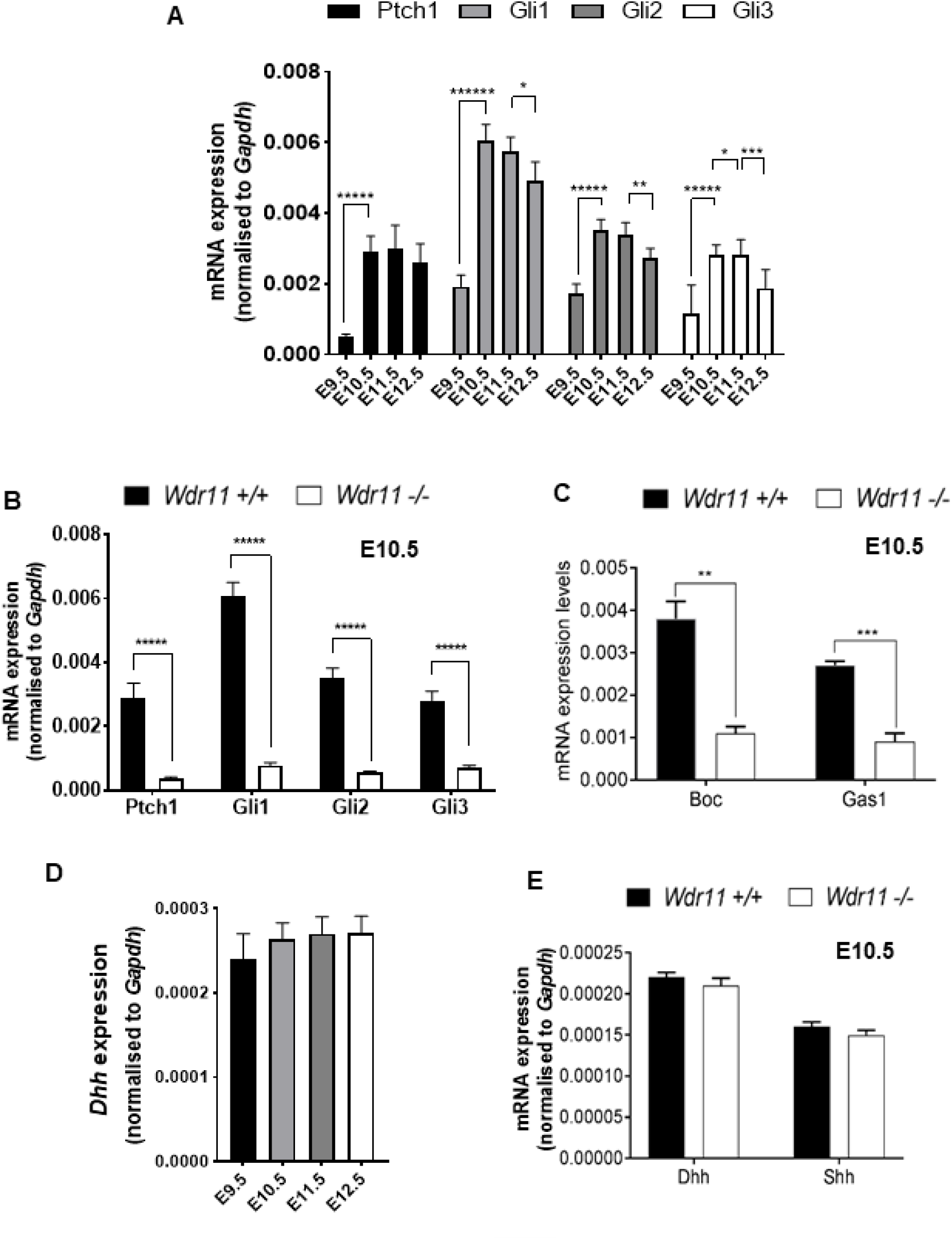
Expression of Hh pathway genes in the PGC migratory niche is disrupted by Wdr11 KO. **(A)** Expression of Ptch1, Gli1, Gli2 and Gli3 in the PGC migratory niche was assessed by RT-qPCR of WT mouse embryos at E9.5, E10.5, E11.5 and E12.5. Data were normalised to Gapdh. Means ± SD are shown. Statistical analysis by multiple t-test (number of embryos for each stage, n=5). *P < 0.01, **P < 0.001; ***P < 0.0001; *****P < 0.000001. **(B)** Expression levels of Ptch1, Gli1, Gli2 and Gli3 are significantly reduced in the GR area of Wdr11-deficient embryos compared to WT litter mates at E10.5. Means ± SD are shown. Statistical analysis by multiple t-test (number of embryos for each genotype, n=5); **P = 0.003730; ***P = 0.000045; *****P < 0.000001). **(C)** Expression of Boc and Gas1 are significantly reduced in the GR area of Wdr11-deficient embryos compared to WT litter mates at E10.5. Means ± SD are shown. Statistical analysis by multiple t-test (n=5, number of embryos for each genotype; **P = 0.003730; ***P = 0.000045; *****P < 0.000001). **(D)** Dhh mRNA levels did not show statistically significant difference in GRs of E9.5 – E 12.5 embryos (Welch’s ANOVA test). **(E)** mRNA levels of Dhh and Shh in the GRs at E10.5 was not altered by loss of Wdr11 (P = 0.76; P = 0.29 respectively).

**Figure 8.**
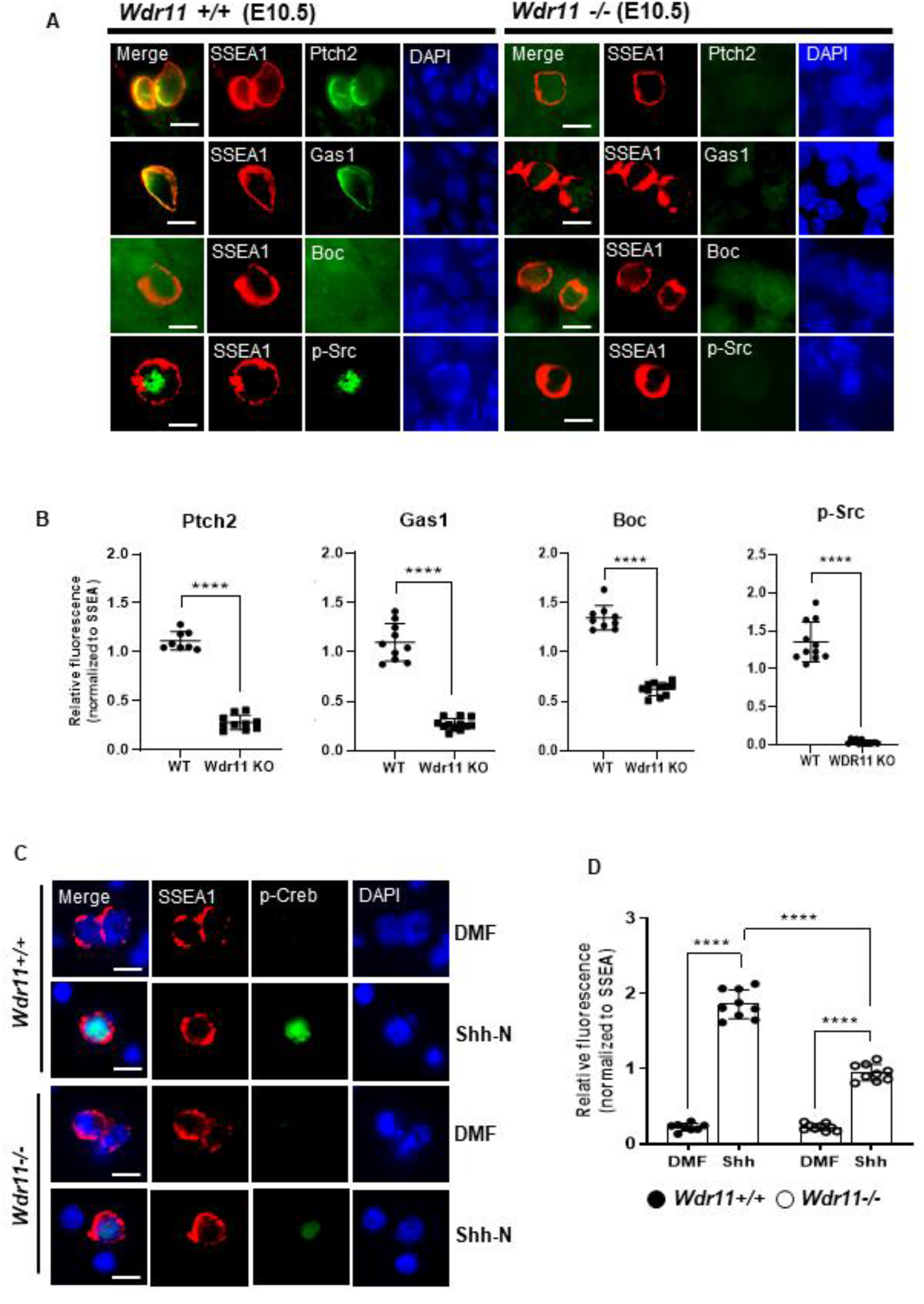
Loss of Hh receptors and p-Src in Wdr11-deficient PGCs. **(A)** Immunofluorescence analyses of Ptch2, Gas1, Boc and p-Src on the GR sections of WT and Wdr11 KO embryos at E10.5. PGCs are labelled by SSEA1 staining. The merged images are shown without DAPI signal to improve the clarity. Scale bar, 10μm. **(B)** The relative fluorescence intensity values of Ptch2, Gas1, Boc and p-Src, that were normalised with the fluorescence intensity values of SSEA1 in each cell. Data are obtained from WT (n=8), KO (n=10) for Ptch2; WT (n=10), KO (n=12) for Gas1; WT (n=9), KO (n=11) for Boc; WT (n=11), KO (n=12) for p-Src. Error bars represent means ± SD. Statistical analysis by unpaired t-test with Welch’s correction. ****P < 0.0001. **(C)** PGCs in the GR primary cultures generated from E10.5 embryos of WT and Wdr11 KO were analysed after immunofluorescence co-staining of p-Creb and SSEA1. Cells plated on 0.1% gelatin-coated cover slips were treated with solvent dimethyl formamide (DMF) or recombinant Shh protein (Shh-N) for 10 minutes. Representative images are shown. Scale bar, 10μm. **(D)** The relative fluorescence intensity values of p-Creb normalized with the intensity values of SSEA1 observed in each PGC were compared in each of the genotype group, with or without Shh-N treatment for 10 minutes. Data are obtained from DMF (n=8) and Shh-N (n=9) for WT; DMF (n=9) and Shh-N (n=9) for Wdr11 KO. Error bars represent means ± SD. Statistical analysis by unpaired t-test with Welch’s correction. ****P < 0.0001.

### Wdr11 KO affects non-canonical Hh signalling in PGCs

Loss of primary cilia cannot explain the reduced growth and migration of PGCs in Wdr11 KO because PGCs are naturally unciliated and can receive Hh signalling through cilia-independent mechanisms (20). Thus, additional mechanisms must be involved. We have recently shown that post-specification migration of PGCs, as observed at E9.5 - E11.5, may be mediated by non-canonical Hh signalling. Locally secreted low concentration of Hh was required for maintenance of the intrinsic motility of PGCs (20). Desert hedgehog (Dhh) has also been associated with genitourinary tract development (39). Our qRT-PCR data showed Dhh was indeed expressed in the PGC niche at a slightly higher level than Shh, although there was no significant changes of Dhh during E9.5 – E12.5 (Fig 7D). Therefore, we investigated if the expression of Shh or Dhh was altered in Wdr11-null GRs. Interestingly, Wdr11 KO did not affect the expression of either Dhh or Shh (Fig 7E). These data demonstrate that the reduced proliferation and motility of PGCs in Wdr11 KO was not due to a reduced expression of Hh ligands themselves. If so, it may be the accessibility or reception of Hh ligand that is defective in Wdr11 KO embryos, preventing the activation of downstream effector pathways.

It was reported that the Hh ligand is cooperatively received by Ptch2 and its co-receptor Gas1, exclusively expressed on PGCs (20). Upon stimulation with Hh ligand, the Ptch2/Gas1 hetero-complex mediated the rapid de-repression of Smo and induced non-canonical Hh signalling within minutes rather than days as required by the canonical Hh signalling associated with Gli transcription factors. This Ptch2/Gas1-dependent signalling did not require translocation of Smo to the primary cilium. Therefore, the unciliated PGCs could still respond to Hh ligands (20). To further define the role of Wdr11 in this context, we investigated the status of these Hh receptors in WT and Wdr11−/− PGCs. Immunofluorescence analyses showed that Ptch2 and Gas1 expression was virtually absent from Wdr11 KO PGCs (Fig 8A and B). Gas1 mRNA level was also markedly reduced (Fig 7C), indicating that Wdr11-defective PGCs may be unable to respond to Hh due to the lack of Hh receptors.

The nonreceptor tyrosine kinase Src is a regulator of cell motility and proliferation and was shown to be activated by phosphorylation in migrating PGCs via Ptch2/Gas1-dependent Hh signalling (20). Our immunofluorescence analyses of GR tissues using phospho-Src antibody demonstrated that Wdr11-null PGCs exhibited significantly reduced activation of Src (Fig 8 A and B). Therefore, defective expression of Ptch2/Gas1 on the Wdr11-null PGCs led to a failed induction of downstream signalling effectors required for motility and proliferation such as p-Src. It has also been demonstrated that Ptch2/Gas1-dependent Hh signalling can elicit a global induction of cAMP signalling and phosphorylation of Creb in the cytoplasm, which is abolished in the absence of either Ptch2 or Gas1 (20). Since activation of Creb has a pivotal role in cell proliferation and motility (40) and agents that increase intracellular cAMP levels such as forskolin are shown to enhance PGC proliferation (41), we sought to determine if Wdr11-deficient PGCs failed to induce p-Creb in response to Hh. To this end, we generated primary cultures of GR tissues and stimulated them with recombinant Shh protein (Shh-N) for 10 minutes, which was shown to induce p-Creb in PGCs but not in the somatic cells (20). Primary GR cultures which were serum starved for 24 hours exhibited very little basal p-Creb. Shh-N induced a significant upregulation of p-Creb in WT PGCs, which was markedly attenuated in Wdr11 KO PGCs (Fig 8C and D). Combined, these data support the notion that Ptch2/Gas1-dependent non-canonical Hh signalling involving Src and Creb is disrupted in Wdr11-null PGCs.

### Effects of Wdr11 mutations in mesenchymal cell proliferation

In an attempt to directly confirm the role of Wdr11 in mesenchymal cell proliferation and to predict the consequences of disease-associated mutations of Wdr11, we employed NIH3T3 cells as a model and engineered a targeted KO of Wdr11 by CRISPR/Cas9-mediated gene editing. In addition, we also introduced two clinically identified missense mutations of WDR11, namely MT and RC variants. The MT mutation (c.1610C>T; p.Pro537Leu) was originally found in two brothers with delayed puberty and childhood obesity (12). The RC mutation (c.1783T>A; p.Trp595Arg) was found in a 61-year old male patient with high grade clear-cell renal cell carcinoma (42). We also generated a targeted KO of IFT88, a gene critically required for ciliogenesis, disruption of which is known to cause defective cilia formation and function (43). The specific mutations and targeted KO were confirmed by both Sanger sequencing of genomic DNA and Western blotting of the endogenous proteins (Fig 9A). These genetic manipulations of NIH3T3 cells did not affect the gross cell morphology and the general cytoskeletal architecture (Fig 9B). However, when we examined the status of primary cilia, mutant cells showed a significant reduction in the cilia length compared to the WT (Fig 9C), except Wdr11-RC mutant which still maintained a comparable cilia length. Notably, ciliation frequency of Wdr11 mutants did not differ significantly from that of WT cells, while Ift88 KO caused a severe reduction in both cilia length and frequency (Fig 9C). We then asked whether these mutations altered cell proliferation by recording cell counts in normal growth medium over 3 days. Compared to the WT, cells with Wdr11 KO showed a severely attenuated growth, while those expressing Wdr11-MT and IFT88 KO showed a relatively moderate inhibition. Interestingly, the highest rate of proliferation was observed in Wdr11-RC mutant which had least affected cilia (Fig 10A), potentially implying pro-mitogenic effects associated with malignant cancers. The correlation between the proliferative capacity and the degree of cilia shortening suggests that cilia-dependent canonical Hh signalling may regulate the proliferation of these cells. We speculate that a similar mechanism may underlie the reduced proliferation of somatic cells in the Wdr11-deficient GR exhibiting defective canonical Hh signalling (Fig 7).

**Figure 9.**
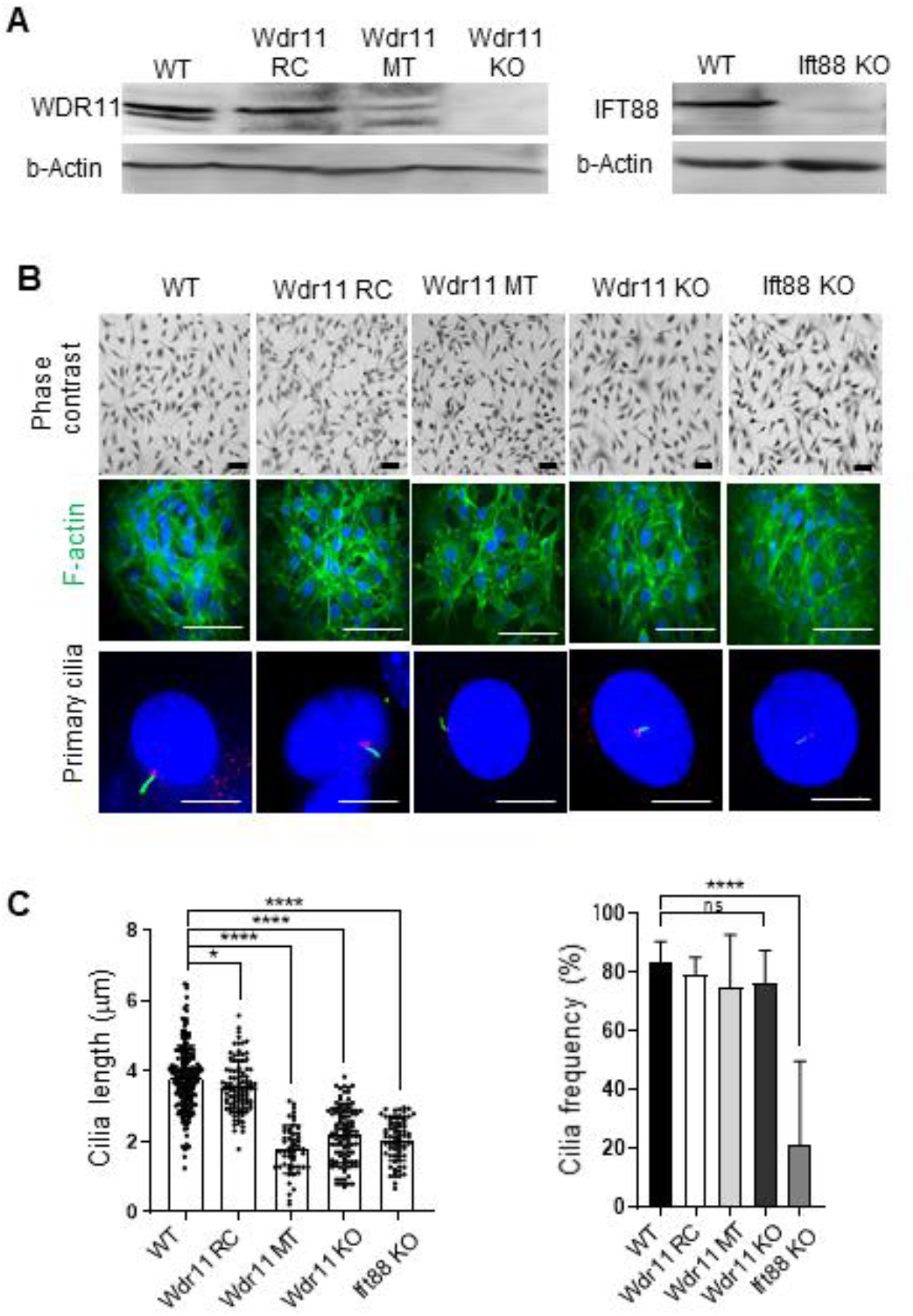
Characterisation of NIH3T3 cells after CRISPR/Cas9-mediated mutagenesis. **(A)** Western blot analyses of NIH3T3 cells with targeted gene editing confirmed the absence of endogenous proteins after KO. The missense variants Wdr11-RC and Wdr11-MT still produced the full-length proteins although at reduced levels. All variants were also confirmed by direct Sanger sequencing. **(B)** The phase contrast microscope imaging (scale bar 100μm) and F-actin phalloidin staining (scale bar 100μm) showed that the gross cell morphology and cytoskeletal organisation of different NIH3T3-CRISPR/Cas9 cells were not altered significantly. The primary cilium (scale bar 10μm) was visualised with Arl13B labelling for axoneme (green) and gamma-tubulin labelling for basal body (red). Representative images are shown. **(C)** NIH3T3 CRISRP/Cas9 cells plated onto the glass cover slips coated with 0.001% poly-L-lysine in PBS were incubated in serum-free medium for 24 hours to induce primary cilia formation. The length of the cilia axoneme was measured from WT (n=186), Wdr11-RC (n=98), Wdr11-MT (n=52), Wdr11 KO (n=101) and Ift88 KO (n=68). The ciliation frequency was assessed by counting the total number of nuclei and cilia in the random fields of cells from WT (n=11), Wdr11-RC (n=9), Wdr11-MT (n=10), Wdr11 KO (n=9) and Ift88 KO (n=16). Error bars represent means ± SD after unpaired t-test with Welch’s correction. *P < 0.01, ****P < 0.0001, ns (non-significant).

**Figure 10.**
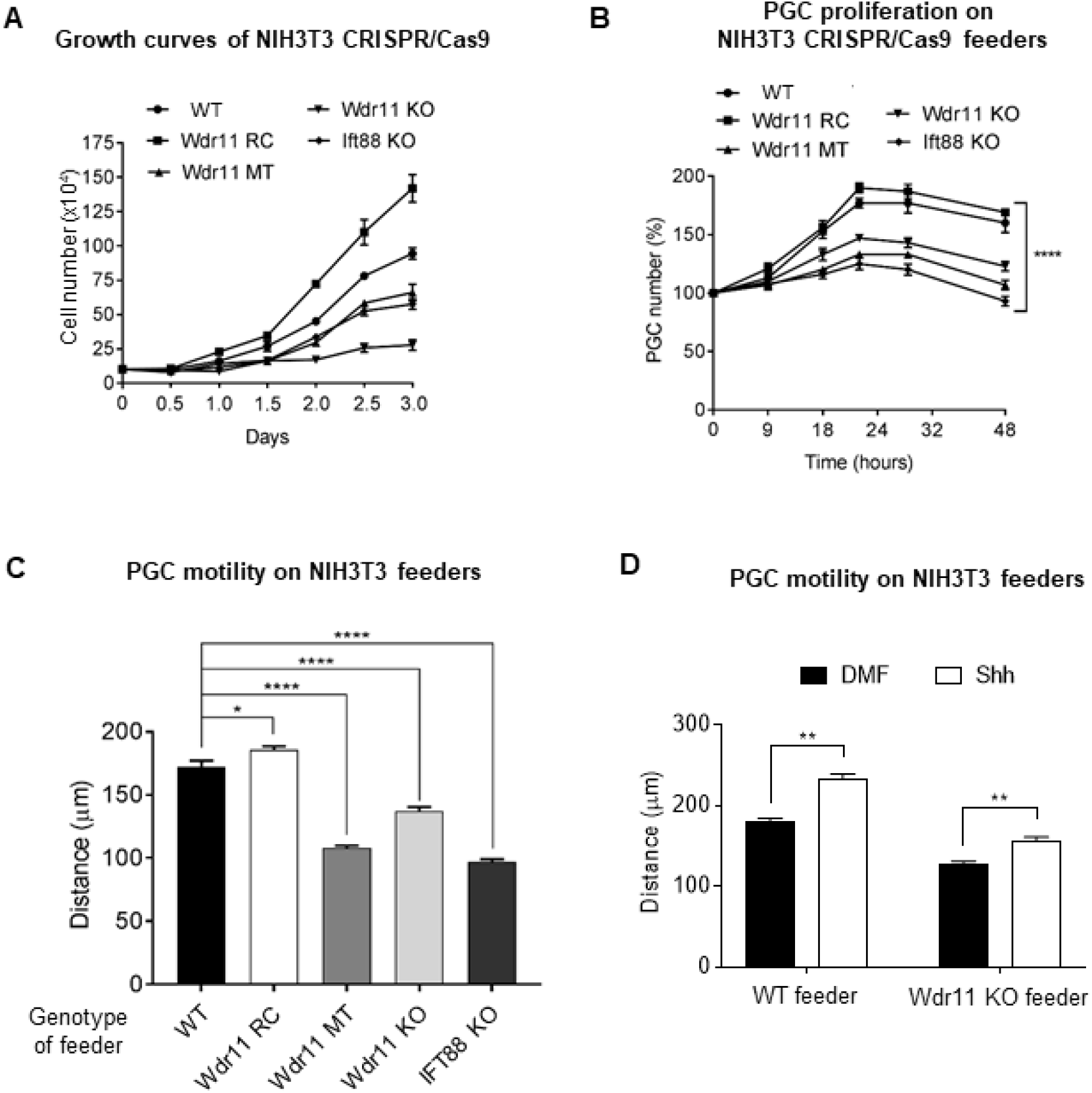
Effects of Wdr11 mutations on primary PGC co-cultures. **(A)** The mitotic effects of various mutations introduced in NIH3T3 cells are shown as growth curves. Cells were plated at 2×10^6^ cells per 10 cm dish in the normal growth medium and counted at the time points indicated. **(B)** Proliferation of PGCs cultured on different NIH3T3 CRISPR/Cas9 feeder cell layers. The growth curves were generated by counting GFP-positive cells from 10 random fields at the time points indicated. The percentage was calculated from the total cell count at 0 hours. **(C)** Intrinsic random motility of PGCs cultured on different NIH3T3 CRISPR/Cas9 feeder cell layers assessed by live time-lapse imaging (see Supplementary Movies 3 - 7). The average of accumulated moving distance of 20 GFP-positive cells in random fields of view tracked for 16 hours in 3 biologically independent experiments are shown. Error bars represent means ± SD after unpaired t-test with Welch’s correction. *P < 0.01, ****P < 0.0001. **(D)** Effects of Shh-N on the motility of PGCs cultured on WT and Wdr11 KO feeder cell layers (see Supplementary Movies 8 - 11). The average of accumulated moving distance of 20 GFP-positive cells in random fields of view tracked for 16 hours in 3 biologically independent experiments are shown. Error bars represent means ± SD after unpaired t-test with Welch’s correction. PGCs on WT feeder were increased by 30.2±0.6% upon Shh-N treatment, while those on Wdr11 KO feeder were increased by 22.9±0.7%.

### Effects of Wdr11 mutations in proliferation and motility of PGCs

PGCs depend on the neighbouring somatic cells for survival and expansion (44–46). It is shown that isolated mouse PGCs can be cultured on feeder cell monolayer treated with Mitomycin-C (47). In such conditions, the feeder cells are not proliferating but physiologically alive, producing soluble factors and surface molecules necessary for stimulating PGC proliferation and motility while preventing apoptosis. To explore the consequences of Wdr11 mutations in the somatic cells which essentially govern the expansion and migration of PGCs, we established a PGC co-culture system where single-cell suspensions of GR tissues were seeded onto NIH3T3-CRISPR/Cas9 feeders expressing different mutations Wdr11 and Ift88. Growth curves of PGCs, generated by counting GFP-positive cells in the co-cultures over 48 hours period, indicated that the mutant feeders expressing Wdr11-MT, Wdr11 KO or Ift88 KO caused a significant reduction in PGC proliferation compared to the WT feeder (Fig 10B). On the contrary, Wdr11-RC feeder supported PGC proliferation almost as effectively as the WT feeder (Fig 10B). These results reinforce the notion that the property of somatic feeder cells can influence the growth of the PGCs.

Next, we investigated the impact of somatic mutations of Wdr11 on the motile capacity of PGCs. To this end, we analysed the random motility of isolated PGCs by time-lapse imaging of the co-cultures seeded on different feeders (Supplementary Movies 3-7). Analyses of the accumulated distance over 10 hours demonstrated that PGCs cultured on WT and Wdr11-RC mutant feeder maintained a similar level of intrinsic motility but PGCs cultured on other mutant feeders had a significantly decreased motility (Fig 10C). This result also validates our slice culture experiment (Fig 3), confirming that the altered PGC migration in Wdr11-null embryo was not simply a consequence of morphological changes in the growing embryo, because even when the PGCs and their neighbouring somatic cells were dispersed and cultured as a monolayer in a dish, the effects of Wdr11 mutant feeders on the PGC motility were clearly demonstrable. Therefore, defective cilia on feeder cells caused by loss-of-function mutations of Wdr11 or Ift88 may have significant impacts on PGC proliferation and migration.

### Shh can rescue defective motility of PGCs

We previously reported that treatment of Hh agonists (Purmorphamine or recombinant Shh-N) increased the motility of PGCs and CRISPR/Cas9-mediated deletion of Wdr11 in NIH3T3 cells abolished the accumulation of Shh in the conditioned medium (20). If so, it is possible that the reduced motility of PGCs cultured on Wdr11 KO feeders might be due to an insufficient supply of Hh ligand or Hh signalling molecules by the feeders. Therefore, we investigated if an addition of exogenous Hh ligand could rescue the reduced PGC motility on Wdr11 KO feeders. The motility of PGCs cultured on Wdr11 KO feeder was markedly lower than those cultured on WT feeders (Fig 10D). However, after Shh-N treatment, it was increased to a level comparable to that of WT feeders. This finding is in line with the idea that Wdr11-null cells with defective cilia may be unable to produce Hh signalling molecules required for PGC motility, which can be partly rescued by addition of exogenous Shh-N (Fig 10D). Combined, these results suggest that the loss of Wdr11 which disrupts the function of primary cilia on the somatic cells affects the behaviour of PGCs, potentially due to a failed provision of Hh ligand necessary to induce the non-canonical signalling required for the migration and proliferation of PGCs.

## DISCUSSION

PGCs migrate independently of one another mainly guided by their interaction with the environment. PGCs also express markers typical of embryonic stem cells including OCT4, NANOG and SOX2 (48). The fact that PGCs and their surrounding somatic cells originated from different niches and the unique property of PGCs naturally lacking primary cilia makes a fascinating model system where cilia-dependent and -independent Hh signalling pathways concurrently regulate cell behaviour in different context. We propose that canonical and non-canonical Hh signalling are differentially involved in the development of somatic and germ cell components of the gonads, and both pathways are each required, in parallel, for normal PGC development. How PGCs receive and interpret these different signals is not fully understood yet, but paracrine secretion or delivery of the Hh signalling component from the primary cilia may be involved. Although we did not specifically address this question in current paper, shedding of ciliary tips or release of ectosomes containing Hh signalling molecules, synchronised with cell cycle, has been reported (49–51). Using the PGC co-culture system, we demonstrated that primary cilia are not only important for the growth of soma itself but also critical for the proliferation and migration of PGCs growing on them. The fact that Ift88 KO feeder also showed a significant impairment in supporting PGCs suggests that it is not just a specific effect by Wdr11, but may be applicable to other ciliopathy genes. We also established the genotype-phenotype correlations of clinically identified mutations of Wdr11, providing new insights for the pathogenesis of Wdr11-associated disorders such as CHH/KS (potential loss of function) and renal carcinomas (potential gain of function).

Defective migration and proliferation of PGCs in Wdr11 mutants cannot simply be a secondary manifestation of primary cilia defects leading to a multitude of developmental signalling failures, because we found that other key regulators of PGCs were not severely affected by Wdr11 KO (Fig 5). We have recently shown that the local Hh signalling is required for the migration of post-specification PGCs. Treatment with recombinant Shh or a Smo agonist could enhance PGCs’ intrinsic motility, while Hh antagonists such as cyclopamine and vismodegib inhibited it without affecting directionality (20). The unciliated PGCs rely on Ptch2/Gas1-dependent non-canonical Hh signalling pathway mediated by p-Src and p-Creb. Conversely, Ptch1/Boc-dependent canonical Hh signalling is likely responsible for the maintenance of the surrounding mesenchymal cell proliferation via Gli transcription factors. Our study provides further evidence for the requirement of these signalling pathways in PGC development, which is disrupted by Wdr11 KO.

At least two lines of evidence suggest that the migrations of developing GnRH neurones and PGCs are linked or mediated by common signalling pathways. First, chemokine SDF-1 and its receptor CXCR4, which have an established role in directed migration of PGCs, can also regulate GnRH neuronal migration (18). SDF-1 is expressed in the nasal mesenchyme, whereas CXCR4 is localised in migrating GnRH neurons and olfactory/vomeronasal nerve axons. *Cxcr4*-deficient mice contained significantly decreased numbers of GnRH neurones accompanied by defective migration. Secondly, Fibroblast Growth Factor (FGF) signalling pathway, a well-established KS/CHH-associated signalling pathway involving genes such as FGFR1, FGF8 and Heparan sulfate 6-O-sulfotransferase 1 (52–54), is also important in development of PGCs (55). PGCs express two FGF receptors, FGFR1-IIIc and FGFR2-IIIb. FGF2, the ligand for FGFR1-IIIc, modulates PGC motility whereas FGF7, the ligand for FGFR2-IIIb, affects PGC proliferation (55). Importantly, FGFRs are also shown to localise to primary cilia, affecting the length and function of primary cilia (56). Loss of FGFR1 or its FGF ligands resulted in shorter cilia in zebrafish and *Xenopus* (57). Based on these findings we speculate that KS/CHH might be a ciliopathy. KS/CHH are traditionally considered as a 'secondary' hypogonadism caused by defective development and function of GnRH neurones. Our data suggest that 'primary' hypogonadism (defects within the gonads) may also contribute. In a study, 26% of KS/CHH male patients did not respond to GnRH therapy (i.e. normalisation of testosterone levels, testes volume and spermatogenesis), suggesting that primary testicular defects may be involved (4). Nonetheless, these atypical responders were still considered as secondary (i.e. hypo-gonadotrophic) hypogonadism because GnRH administration could still increase LH/FSH levels. So far, up to 56 genes are reported to be associated with KS/CHH. Many of these genes are broadly expressed in the HPG axis; thus, the impact of the mutations may not be limited to hypothalamic GnRH neurons but produce more than one primary defect within the HPG axis. Those KS/CHH patients who do not respond to gonadotrophin therapies may suggest the possibility of primary hypogonadism, resulting from being born with significantly reduced numbers of germ cells. Investigation of the effects of other KS/CHH-associated genes in PGC migration may provide further evidence.

Here we report a previously undescribed role for Wdr11 in development of the germ line with direct consequences in PGC development. Loss of *Wdr11* resulted in defective cilia and disrupted GnRH neuronal migration in mouse and zebrafish in vivo. ShRNA-mediated knockdown of WDR11 in human GnRH neuronal cells caused defective cilia, which was partially rescued by Hh agonist in vitro (12). Several members of the WDR protein family have been shown to play important roles in ciliogenesis. WDR11 is a multi-functional adaptor protein involved in cargo trafficking to the trans-Golgi network (58). Other studies have implicated WDR11 as a part of adaptor complexes regulating ciliogenesis (59) and autophagy (60). We speculate that WDR11 is involved in the assembly and reabsorption of the primary cilium via endosome trafficking and targeted protein degradation at the ciliary base. The clinically identified mutations of WDR11 are mostly found on the surface of the protein, thus are likely to interfere with the interactions of other binding partners (7).

Hypogonadotrophic hypogonadism and infertility with or without anosmia are part of the clinical features of ciliopathies such as Bardet-Biedl syndrome (61), but in most cases, ciliopathy-related infertility was considered to be caused by the motile cilia defects affecting sperm flagella and oviduct epithelium. Our data show that non-motile primary cilium-dependent mechanisms also play an important role. Primary cilia-dependent Hh signalling is required for the proliferation and migration of PGCs that populate the foetal gonads, the lack of which could account for the germ cell insufficiency at birth, potentially leading to subfertility or infertility. Although several genes ensuring proper development of germ cells have been identified (14), the link with KS/CHH has not been established. KS/CHH patients show small testes and ovarian insufficiency, raising the possibility that normal gametogenesis might be still possible with the remaining germ cells, since loss of Wdr11 does not appear to affect the specification or fate determination of PGCs. Understanding the role of Wdr11 in later stages of reproductive development including gonadogenesis and steroidogenic cell differentiation will require further studies.

## MATERIALS AND METHODS

### Breeding of transgenic mice

*Stella*^*GFP*^ mice were originally obtained from Azim Surani (Gurdon Institute) (31) and maintained in a C57BL/6 background as described (20). The Wdr11 knockout mouse (International Gene Trap Consortium Ayu21-KBW205) was generated at the Institute of Resource Development and Analysis, Kumamoto University in Japan (12). To establish the *Stella*^*GFP+/+*^;*Wdr11*^+/−^ hybrid line, the homozygote *Stella*^*GFP*^ mice were crossed with the heterozygote Wdr11 mice. The noon copulation plug was counted as embryonic day 0.5 after timed mating. All experiments were conducted in accordance with the Animals (Scientific Procedures) Act 1986 in the Biological Research Facility at St. George’s, University of London (PPL 70/8512) according to approved institutional guidelines and protocols.

### Mouse genotyping

The genotypes of the parent mice and their litters were verified by performing PCR and qPCR analyses of the genomic DNA. The copy number of the GFP allele was determined quantitatively by qPCR to confirm the genotypes. To perform the relative quantification, the crossing point (Cp) value of the target gene (*GFP*) was normalised to the Cp of the reference gene (*β-tubulin*), based on which the Relative Copy Number Ratios (RCNR) were generated. Copy Number Variation (CNV) was calculated by 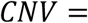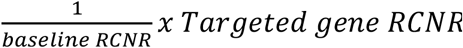. The rounded CNV value of 1 was considered to indicate a heterozygote and 2 a homozygote. A CNV value of 0 indicated a WT mouse (i.e. no GFP). The presence of the *Stella*^*GFP*^ allele was further confirmed by test-breeding of randomly selected homozygous litters with WT mice, followed by PCR amplification of GFP. Primers used for genotyping analyses are shown in Supplementary Information.

### qPCR and RT-PCR

For RT-PCR analyses, mouse tissues were harvested and homogenized before total RNA was extracted using an RNeasy Mini Kit according to the manufacturer’s protocol. First-strand complementary DNA (cDNA) was synthesized using oligo(dT) primers and the Precision nanoScript2 Reverse Transcription Kit (Primer Design). Quantitative real-time PCR was performed using the Maxima^®^ SYBR green qPCR master mix (Thermo Fisher Scientific) in a Light Cycler 2.0 instrument (Roche). The crossing point (Cp) values were obtained by LightCycler^®^ Version 4.1 software (Roche). Cp values were analysed using the 2^−ΔΔCT^ method normalised to *Gapdh*. All primers used are provided in Supplementary Information.

### Immunohistochemistry

Embryos were fixed in 4% paraformaldehyde prior to paraffin embedding. Sections cut at 6 μm thickness were de-paraffinised with Histoclear (National Diagnostics) and rehydrated in PBS. For β-galactosidase detection, whole-mount embryos were fixed with X-gal Fix buffer (0.2% glutaraldehyde, 2% paraformaldehyde, 5mM EGTA, 2mM MgCl2 in PBS pH 7.4) for 1 hr at 4°C, washed in PBS and then incubated overnight at 37°C in X-gal solution (1mg/ml X-gal, 2mM MgCl_2_, 5mM K3Fe(CN)6 in PBS at pH 7.4). After washing in PBS and paraffin-embedding, samples were sectioned at 12µm-thickness and counterstained with eosin. For alkaline phosphatase (ALP) staining, embryo sections were stained with BCIP-NBT (Roche) in ALP buffer at 4°C. Images of embryo sections were analysed by Zeiss Axioplan 2 Upright.

### Immunofluorescence

Serial sections of dissected embryos at 5-7μm thickness were deparaffinized, rehydrated and washed in PBS. Following antigen retrieval in sodium citrate buffer (10 mM sodium citrate, 0.05% Tween 20, pH 6.0), sections were blocked with 10% goat serum in 0.5% Triton-X PBS for 1 hour at room temperature and then incubated overnight at 4 °C with primary antibodies diluted in 10% goat serum in 0.5% Tween in PBS. After washing, samples were incubated with fluorescence-labelled secondary antibodies at 1:500 dilution and counterstained with DAPI before mounting in Mowiol. For immunofluorescence analyses of cultured cells, cells were plated on glass coverslips, fixed with 4% PFA, permeabilized with 0.2% Triton X‑100 in PBS, and incubated in blocking buffer (2% heat-inactivated goat serum, 0.2% Triton X‑100 in PBS) before probing with primary antibodies diluted in blocking buffer. After washing, secondary antibodies were added along with DAPI. Fluorescence microscopy was performed using a Zeiss Axiovert 200M Upright microscope and analysed by ImageJ software (http://rsbweb.nih.gov/ij/).

### Imaging of primary cilia and F-actin

Cultured cells on glass cover slips were serum-starved for 18-24 hours before fixing to induce ciliogenesis. Embryo sections were prepared as above. Samples were analysed by immunofluorescence staining with anti-Arl13B antibody that visualises the cilia axoneme or anti-gamma-tubulin antibody that visualises the basal body. To generate ciliation frequency values, the total number of cilia and nuclei were counted from the maximum intensity projection images of each channel manually. The length of cilia was assessed in random fields of cells after Arl13b staining by measuring the maximum projection using ImageJ. To generate the 3D imaging of GR section, 3D volume rendering of the image stacks was performed in ImageJ software using the volume viewer plugin. For F-actin staining, Alexa Fluor 488-conjugated phalloidin (Invitrogen, LSA12379) was used.

### Embryo slice culture and live imaging

Embryo slice organ culture and filming was performed as previously described (20). Briefly, transverse sections of E10.5 embryos were cultured in Hepes-buffered DMEM/F-12 medium with 0.04% lipid-free BSA and 100U/ml penicillin/streptomycin. A single optical section was captured every 15 min for approximately 10 hrs (total 40 frames). The z-stack images were extracted as TIFF files and one stack per time interval was put together using ImageJ to create a movie. Motile behaviour of PGCs was evaluated based on accumulated distance (total cell path travelled), Euclidean distance (the shortest distance between cell start and end points), cell velocity and directionality (the ratio between Euclidean distance and accumulated distance indicating the straightness of the migration path), using the Chemotaxis and Migration Tool 2.0 plug-in software (Ibidi GmbH). Velocity measurements were generated for each time interval by using the formula V = [sqrt (dx^2^ + dy^2^)](p)/0.25h, where dx is the change in the x-axis, dy is the change in the y-axis, and p is the pixel size in μm. The velocities of all the tracked cells were averaged to obtain an overall mean velocity for each embryo slice/movie. Tracking was performed only on those PGCs that remained in focus and viable for the entire duration of filming. Ectopic PGCs localised in the mesentery and hindgut were not analysed as they tend to disintegrate during filming.

### Genital ridge primary culture and live imaging

Dissected GR tissues of E10.5 embryos were digested in 0.25% trypsin, passed through a 0.4μm cell strainer and suspended in DMEM/L-15 medium supplemented with 20% knockout serum replacement (Invitrogen), 2mM L-glutamine, 0.1mM non-essential amino acids and 0.1mM 2-mercaptoethanol (Sigma-Aldrich), before being plated onto 0.1% gelatin-coated cover slips. Cells were incubated in 0.5% serum-containing media before treatment with 200 ng/mL recombinant Shh N-terminal peptide (R&D Systems, 1314-SH) diluted in dimethyl formamide (DMF).

For PGC co-cultures with feeder layers, single cell suspensions generated from dissected GR tissues were plated onto the NIH3T3 feeder layer pre-treated with Mitomycin-C (5μg/ml). Motile behaviors of PGCs were measured by time-lapse imaging of GFP-positive cells captured every 15 minutes for 10 hours. Live imaging was performed using Nikon A1R laser scanning confocal microscope in a humidified 5% CO2 chamber at 37.0±0.5°C. Random motility of PGC was analyzed using the Chemotaxis and Migration Tool 2.0 plug-in software (Ibidi GmbH).

### NIH3T3 cell culture and CRISPR/Cas9

NIH 3T3 cells (American Type Culture Collection, Manassas, VA) were routinely cultured in DMEM with 2mM L-glutamine and 100μg/ml penicillin/streptomycin (Sigma-Aldrich), supplemented with 10% newborn calf serum (NCS). For growth curve analyses, NIH3T3 cells were plated at 2×10^6^ cells per 10 cm dish in the growth medium and total cell counts were assessed every 12 hours. NIH3T3 cells with targeted editing of Wdr11 and IFT88 were generated using CRISPR/Cas9 approach. Briefly, sgRNAs designed using the CRISPR Design Tool (http://crispr.mit.edu) were cloned into pSpCas9(BB)-2A-Puro (Addgene #48139) and transfected using Polyfect (Promega). To isolate single-cell clones, transfected cells were plated in 96-well plates. After selection in puromycin (Cambridge Bioscience), positive clones were confirmed by Sanger sequencing and western blot. The sequences of the sgRNA and primers used are provided in Supplementary Information.

### Apoptosis and proliferation analyses

SSEA1-positive PGCs with co-localised staining of phosphohistone-H3 and cleaved caspase-3 were counted from every other sections of the entire length of the gonadal ridge of E10.5 embryos from each genotype. DAPI was used to determine the total number of cells. DAPI-positive cells negative for SSEA1 were counted as somatic cells. The PGC growth curves were generated by counting GFP-positive cells from 10 random fields of GR primary cultures plated on NIH3T3 feeder layer at 0, 9, 18, 24, 32 and 48 hours after plating. The percentage fold was calculated from the total cell count at 0 hours.

The images were captured using an Olympus IX70 inverted microscope (Hamamatsu C4742–95, Hamamatsu, Japan).

### Western blot

Total protein extracted in a lysis buffer (50mM HEPES, 150mM NaCl, 10% glycerol, 1% Nonidet P-40, and 1mM EDTA) containing protease/phosphatase inhibitors (Sigma-Aldrich) was separated by SDS-PAGE and transferred onto Hybond-ECL membrane (Amersham) before being probed with primary antibodies diluted in blocking buffer (5% skim milk in TBS with 0.05% Tween 20 (TBST)). After washing in TBST, membrane was incubated with horseradish peroxidase-conjugated secondary antibodies before analyses by enhanced chemiluminescence (GE Healthcare). B-Actin was used as loading control.

### Antibodies

Primary antibodies used were against GFP (Rabbit IgG, 1:200, Abcam, ab290), SSEA1 (Mouse IgG, 1:200, Developmental Studies Hybridoma Bank, MC-480), Stella (Rabbit IgG, 1:200, Abcam, ab19878), phospho-histone H3 (Rabbit IgG, 1:500, Millipore, 06-570), cleaved-caspase 3 (Rabbit IgG, 1:200, Cell Signalling, 9661), Arl13B (Rabbit IgG, 1:1000, Proteintech, 17711‐1‐AP), gamma‐tubulin (mouse IgG, 1:1000, Sigma T6557), phospho-Src (Rabbit IgG, 1:200, Invitrogen, 44-660G), phospho-Creb (Rabbit IgG, 1:200, Cell Signaling, 9198), Wdr11 (rabbit IgG, 1:100, Abcam, ab175256; goat IgG, 1:100, Santa Cruz, sc-163523), IFT88 (rabbit IgG, 1:500, Proteintech, 13967-1-AP) and b-actin (rabbit IgG, 1:500, CST, 4967L). Secondary antibodies, all of which were from Invitrogen Thermo Fisher Scientific and used at 1:5000 dilution, include Alexa Fluor 488 (Goat anti‐ rabbit, A-11008), Alexa Fluor 555 (Goat anti‐mouse, A-21422), Alexa Fluor 568 (Goat anti‐rabbit, A-11011), Alexa Fluor 555 (Goat anti-rabbit, A-27039) and Alexa Fluor 488 (Donkey anti-goat, A-11055).

### Statistical analyses

Statistical analyses were performed using GraphPad Prism 5 (La Jolla, CA, USA). The numbers of independent replicated experiments (n) are indicated in the relevant figure legends where possible. In some experiments where percentage values are indicated, the values were calculated from the average raw data value of the sample divided by the average raw data value of the control. Significance was tested using an unpaired student’s t-test. In experiments where the number of repeated measures was unequal per group, a one-way analysis of variance (ANOVA) was used with Welch’s test.

## Acknowledgements

YJK, JYL and S-HK were supported by Medical Research Council (MRC) grant MR/L020378/1 awarded to S-HK and LCL. JYL and YJK were also supported by St George’s Research Bridging Fund Scheme and Global Educational Trust awarded to S-HK. We thank Chris Wylie for his generous help and advice with the embryo slice culture. We appreciate Gregory Perry for technical assistance with confocal microscopy live imaging.

## Author contributions

Conceptualization: S-HK, LCL, H-GK; Funding acquisition: S-HK, LCL; Supervision: S‐ HK, NAB, DB; Methodology (creation of models and design of methods): YJK, JYL, S-HK, D-WK, PA; Investigation (performing the experiments): YJK, JYL; Original draft writing: S‐HK; Manuscript review and editing: YJK, JYL, H-GK, D-WK, DCB, NAB, LCL and S-HK.

## SUPPLEMENTARY TABLES

**Supplementary Table 1.**
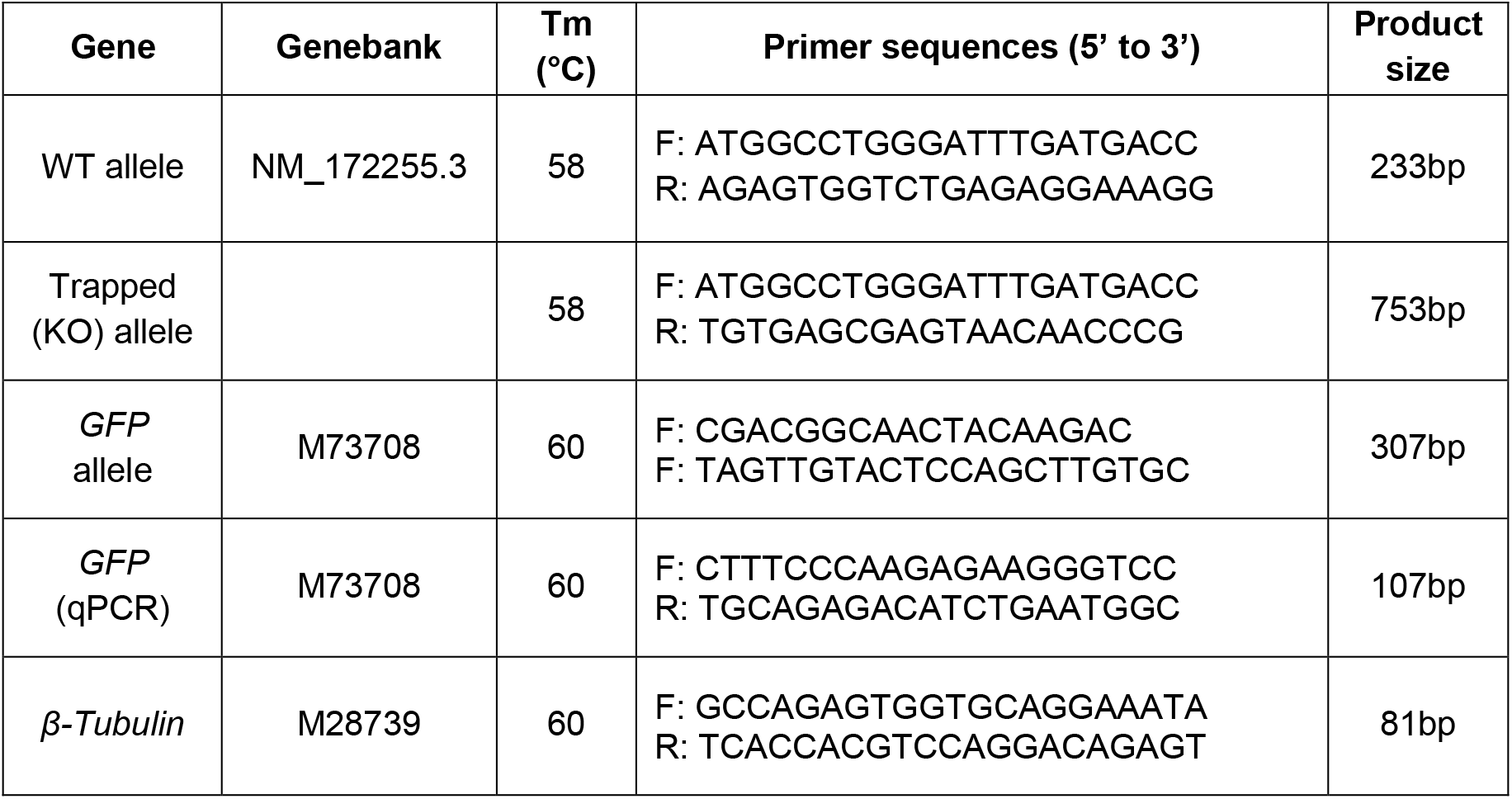
List of primers used for mouse genotyping.

**Supplementary Table 2.**
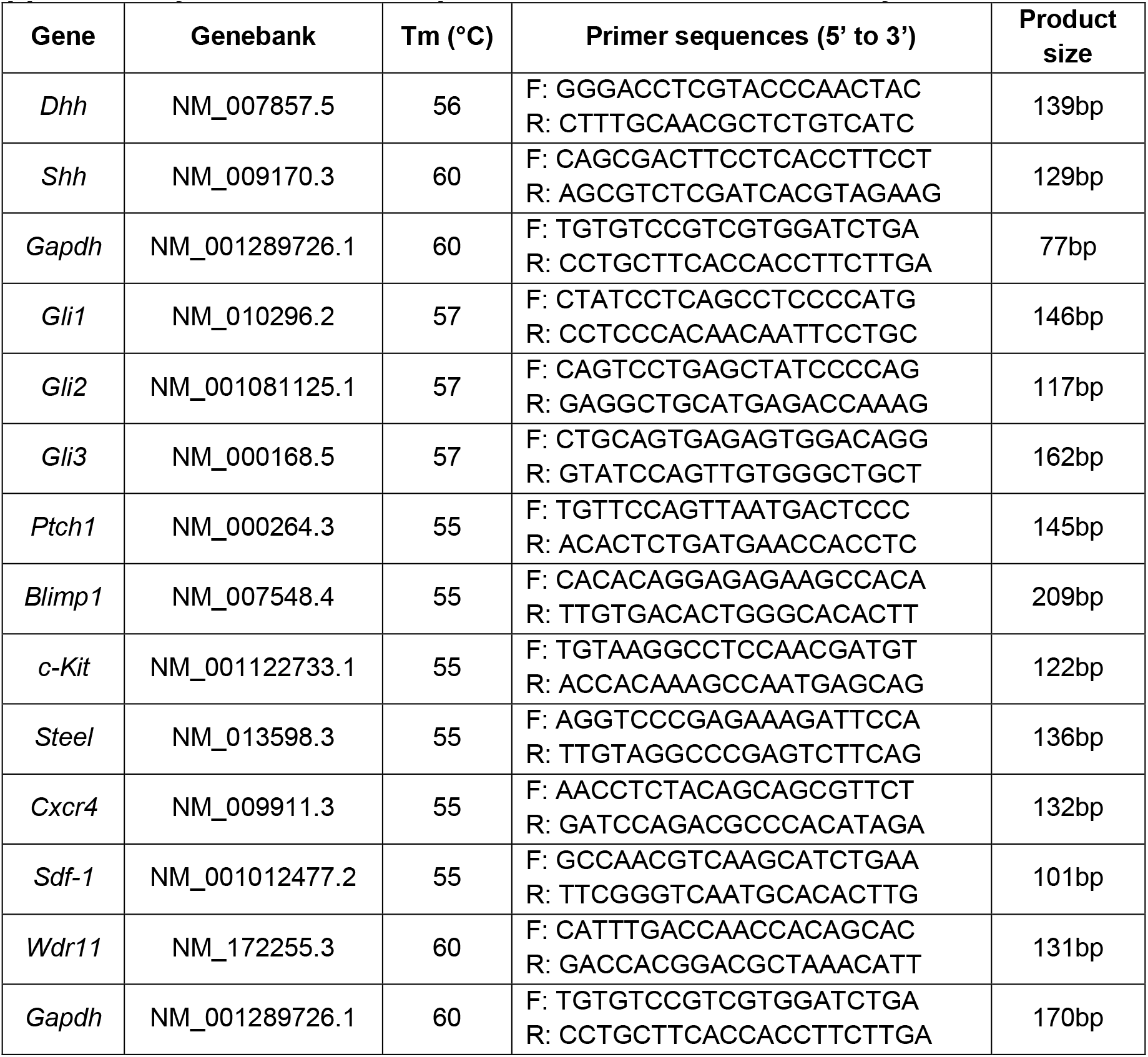
List of primers used for RT-PCR analysis.

**Supplementary Table 3.**
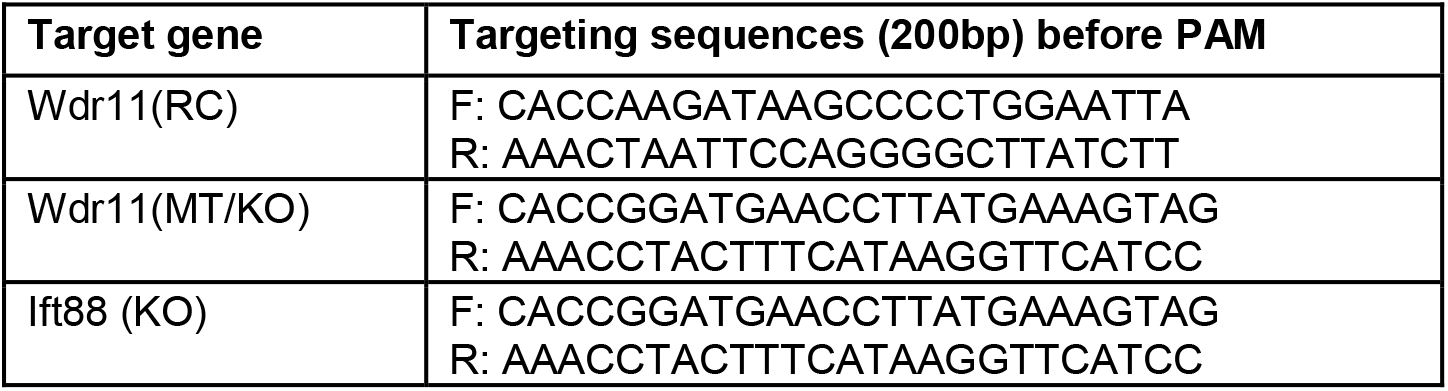
Guide RNAs used for CRISPR/Cas9.

**Supplementary Table 4.**
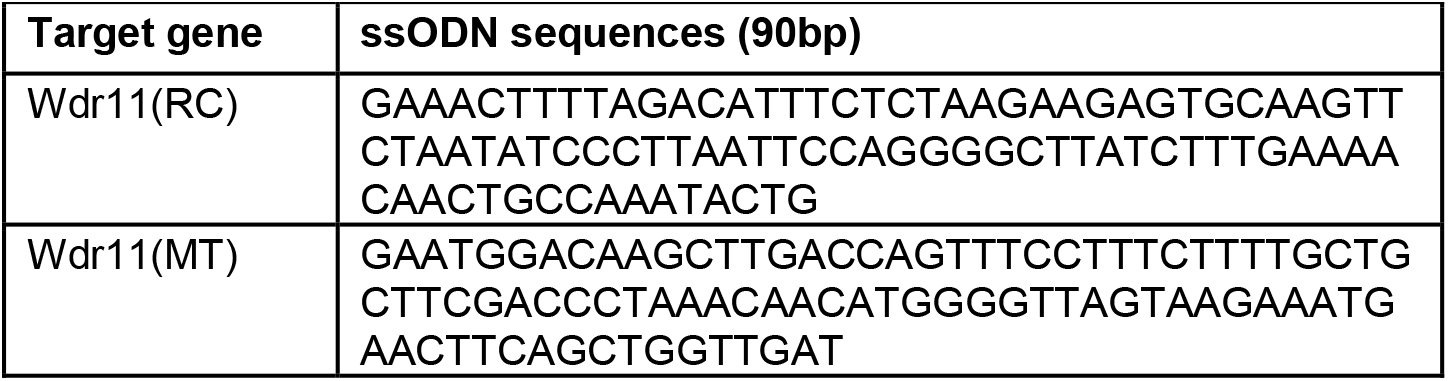
ssODN for generating point mutations.

**Supplementary Table 5.**
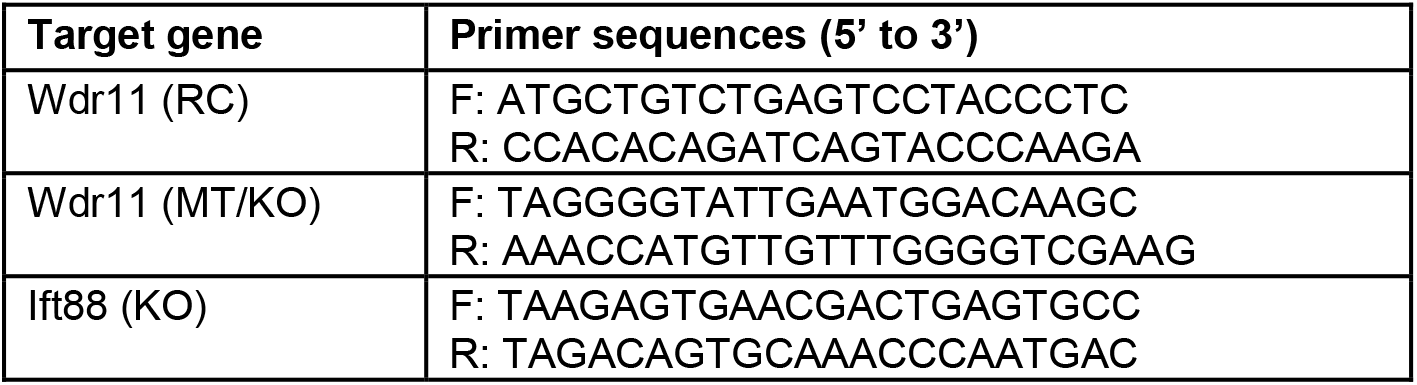
Primers used for validation of gene targeting after CRISPR/Cas9.

## SUPPLEMENTARY MOVIE LEGENDS

**Supplementary Movie 1. PGC migration in E10.5 WT embryo.**

A time-lapse movie of an embryo slice culture from *Stella*^*GFP+/+*^;*Wdr11*^+/+^ mouse as described in the Materials and Methods. PGCs are labelled as green fluorescence.

**Supplementary Movie 2. PGC migration in E10.5 Wdr11 KO embryo.**

A time-lapse movie of an embryo slice culture from *Stella*^*GFP+/+*^;*Wdr11*^−/−^ mouse as described in the Materials and Methods. PGCs are labelled as green fluorescence.

**Supplementary Movie 3. WT genital ridge co-culture on WT feeder.**

A time-lapse movie of E10.5 *Stella*^*GFP+/+*^ mouse GR cells cultured on WT NIH3T3 cell feeder.

**Supplementary Movie 4. WT genital ridge co-culture on Wdr11-RC feeder.**

A time-lapse movie of E10.5 *Stella*^*GFP+/+*^ mouse GR cells cultured on NIH3T3 cell feeder edited for Wdr11-RC mutation.

**Supplementary Movie 5. WT genital ridge co-culture on Wdr11-MT feeder.**

A time-lapse movie of E10.5 *Stella*^*GFP+/+*^ mouse GR cells cultured on NIH3T3 cell feeder edited for Wdr11-MT mutation.

**Supplementary Movie 6. WT genital ridge co-culture on Wdr11 KO feeder.**

A time-lapse movie of E10.5 *Stella*^*GFP+/+*^ mouse GR cells cultured on NIH3T3 cell feeder with Wdr11 KO.

**Supplementary Movie 7. WT genital ridge co-culture on IFT88 KO feeder.**

A time-lapse movie of E10.5 *Stella*^*GFP+/+*^ mouse GR cells cultured on NIH3T3 cell feeder with IFT88 KO.

**Supplementary Movie 8. WT genital ridge co-culture on WT feeder treated with DMF.**

A time-lapse movie of E10.5 *Stella*^*GFP+/+*^ mouse GR cells cultured on WT NIH3T3 cell feeder after treatment with DMF.

**Supplementary Movie 9. WT genital ridge co-culture on WT feeder treated with Shh-N.**

A time-lapse movie of E10.5 *Stella*^*GFP+/+*^ mouse GR cells cultured on WT NIH3T3 cell feeder after treatment with Shh-N.

**Supplementary Movie 100 WT genital ridge co-culture on Wdr11 KO feeder treated with DMF.**

A time-lapse movie of E10.5 *Stella*^*GFP+/+*^ mouse GR cells cultured on NIH3T3 cell feeder with Wdr11 KO after treatment with DMF.

**Supplementary Movie 11. WT genital ridge co-culture on Wdr11 KO feeder treated with Shh-N.**

A time-lapse movie of E10.5 *Stella*^*GFP+/+*^ mouse GR cells cultured on NIH3T3 cell feeder with Wdr11 KO after treatment with Shh-N.

